# Multi-omic screening of invasive GBM cells in engineered biomaterials and patient biopsies reveals targetable transsulfuration pathway alterations

**DOI:** 10.1101/2023.02.23.529575

**Authors:** Joseph H. Garcia, Erin A. Akins, Saket Jain, Kayla J. Wolf, Jason Zhang, Nikita Choudhary, Meeki Lad, Poojan Shukla, Sabraj Gill, Will Carson, Luis Carette, Allison Zheng, Sanjay Kumar, Manish K. Aghi

## Abstract

While the poor prognosis of glioblastoma arises from the invasion of a subset of tumor cells, little is known of the metabolic alterations within these cells that fuel invasion. We integrated spatially addressable hydrogel biomaterial platforms, patient site-directed biopsies, and multi-omics analyses to define metabolic drivers of invasive glioblastoma cells. Metabolomics and lipidomics revealed elevations in the redox buffers cystathionine, hexosylceramides, and glucosyl ceramides in the invasive front of both hydrogel-cultured tumors and patient site-directed biopsies, with immunofluorescence indicating elevated reactive oxygen species (ROS) markers in invasive cells. Transcriptomics confirmed upregulation of ROS-producing and response genes at the invasive front in both hydrogel models and patient tumors. Amongst oncologic ROS, hydrogen peroxide specifically promoted glioblastoma invasion in 3D hydrogel spheroid cultures. A CRISPR metabolic gene screen revealed cystathionine gamma lyase (CTH), which converts cystathionine to the non-essential amino acid cysteine in the transsulfuration pathway, to be essential for glioblastoma invasion. Correspondingly, supplementing CTH knockdown cells with exogenous cysteine rescued invasion. Pharmacologic CTH inhibition suppressed glioblastoma invasion, while CTH knockdown slowed glioblastoma invasion *in vivo*. Our studies highlight the importance of ROS metabolism in invasive glioblastoma cells and support further exploration of the transsulfuration pathway as a mechanistic and therapeutic target.

## MAIN

Glioblastoma (GBM) is the most common and lethal adult brain tumor,^1^ and is biologically characterized by an unparalleled invasive capacity.^2,3^ Current therapeutic strategies are insufficient to control the disease, as reflected by a dismal prognosis including a median survival of less than 15 months from the time of diagnosis.^1^ The current standard of care consists of maximal surgical resection followed by radiation therapy and temozolomide chemotherapy.^4,5^ Unfortunately, the invasiveness of the tumor impedes each of these treatment modalities, as it renders complete surgical resection impossible; leads to the spread of tumor cells outside of the field of radiation; and enables tumor cells to escape the area of MRI enhancement where the blood-brain barrier (BBB) is disrupted into regions outside the enhancement where the BBB is intact, making these invasive cells less accessible to systemic chemotherapy.^2,3^

While numerous hypotheses have been proposed regarding the pathways that regulate GBM invasion, studies thus far have been unsuccessful in identifying specific targetable molecular factors which drive the tumor’s invasive capacity.^2,3^ When investigating biologic processes which might drive invasion in tumors, while it is logical to closely examine the tumor’s cellular metabolism,^6^ as the process of invasion is likely to create a need for GBM cells to shift their metabolic profile in response to the bioenergetic demands of the invasive process and the limited nutrient availability of the surrounding brain,^2,7,8^ the mechanisms enabling GBM cells to fuel their invasive capacity remain understudied.^6,9^

Historically, studies investigating metabolic reprogramming in GBM, as with other cancer types, have focused on glucose metabolism.^7,10^ GBM cells like all cancer cells often metabolize glucose into lactate, even when oxygen is present, a process known as the “Warburg Effect.”^10^ This is hypothesized to allow tumor cells to use glucose-derived carbons for the synthesis of essential cellular ingredients, while still generating sufficient ATP to fuel cellular reactions.^11^ While glucose metabolism is undoubtably important for numerous cellular processes in GBM, recent studies have painted a more complex picture of metabolic reprogramming in this tumor.^10,11^ In addition to a shift towards glycolysis, GBM cells also increase intracellular lipid, amino acid, and nucleotide stores through a variety of molecular mechanisms, including increased extracellular uptake, *de novo* synthesis, and fluxing carbons through numerous biochemical pathways such as the use of glycolysis to provide carbon substrates for the synthesis of nucleic acids.^9,11^ Importantly, these metabolic adaptations respond not only to the tumor’s genotype, but also to features of the surrounding microenvironment such as hypoxia which alters the transcription of metabolic genes.^10^

There is thus a need to apply these advances made in understanding mechanisms of metabolic reprogramming beyond glucose metabolism in GBM to comprehensively define the metabolic alterations needed for GBM invasion. To address this knowledge gap, we employed a multi-omics approach in novel microdissectable biomimetic 3D invasion devices and site-directed biopsies of patient GBMs to define metabolic changes in invasive GBM cells. After validating that our 3D hydrogel platforms adequately and reproducibly reflect the metabolic changes associated with GBM invasion, we then performed a CRISPR screen of metabolic genes and discovered targetable metabolic factors that mediate invasion in this devastating disease.

## RESULTS

### Metabolomics reveals increased cystathionine and other oxidative stress metabolites in invasive GBM cells in 3D hydrogels and patient specimens

To comprehensively analyze the metabolic perturbations in invasive GBM cells, we performed metabolomics analysis of invasive and core GBM cells isolated from 3D hydrogel invasion devices and site-directed patient patients. The 3D hydrogel invasion devices are a modified version of our previously published invasion devices and contain hyaluronic acid (HA) hydrogels decorated with integrin binding peptides (RGD) and crosslinked with protease-cleavable crosslinkers^12^ (**Extended Data Figs. 1a-d**). After long-term culture (28 days), the devices were disassembled, and the hydrogel and cells were microdissected to isolate invasive and non-invasive core cell fractions (n=7; **Extended Data Figs. 1e-f**). Following separation, tumor cells in each of the fractions underwent metabolomic analysis (**Supplementary Table 1**). In parallel, site-directed biopsies were taken from the invasive edge and central core of patient IDH wildtype GBMs (n=5; **Extended Data Fig. 1g**) and subjected to metabolomic analysis (**Supplementary Table 1**). Principal-component analysis confirmed distinct metabolic profiles for invasive and core tumor fractions in both patient tumors and in 3D hydrogels (**Fig. 1a**). Heatmaps were generated to explore the heterogeneity of relative metabolite levels within tumor groups, and to determine whether changes seen between tumor fractions were driven by only a subset of devices or tumors (**Extended Data Fig. 2a**). Volcano plots revealed increased levels of metabolites involved in the response to oxidative stress, including adenosine, spermidine, and nicotinamide, in the invasive fraction of both the 3D hydrogels and site directed patient biopsies (**Fig. 1b**). Most notably, cystathionine, a metabolic precursor to cysteine in the transsulfuration pathway, was upregulated in the invasive fraction of both 3D hydrogels and site directed patient tumor biopsies (**Fig. 1b**) Analysis of the ten most enriched metabolites in the invasive fraction of both 3D hydrogels and patient tumors confirmed cystathionine was in the top two of both groups, the only metabolite to be upregulated to such a high degree in both models (**Fig. 1c**). MetaboAnalyst pathway analysis revealed increased metabolism of glutathione and increased metabolism of several amino acids including cysteine in patient specimens from the GBM invasive edge relative to the GBM core (**Fig. 1d**). Thus, metabolomics confirmed that the metabolomic signature of invasive GBM cells differs from that of core GBM cells, and our analysis suggested that the 3D hydrogel models produced invasive and core GBM cell populations with similar metabolic profiles as those obtained from site-directed patient GBM biopsies.

**Figure 1.**
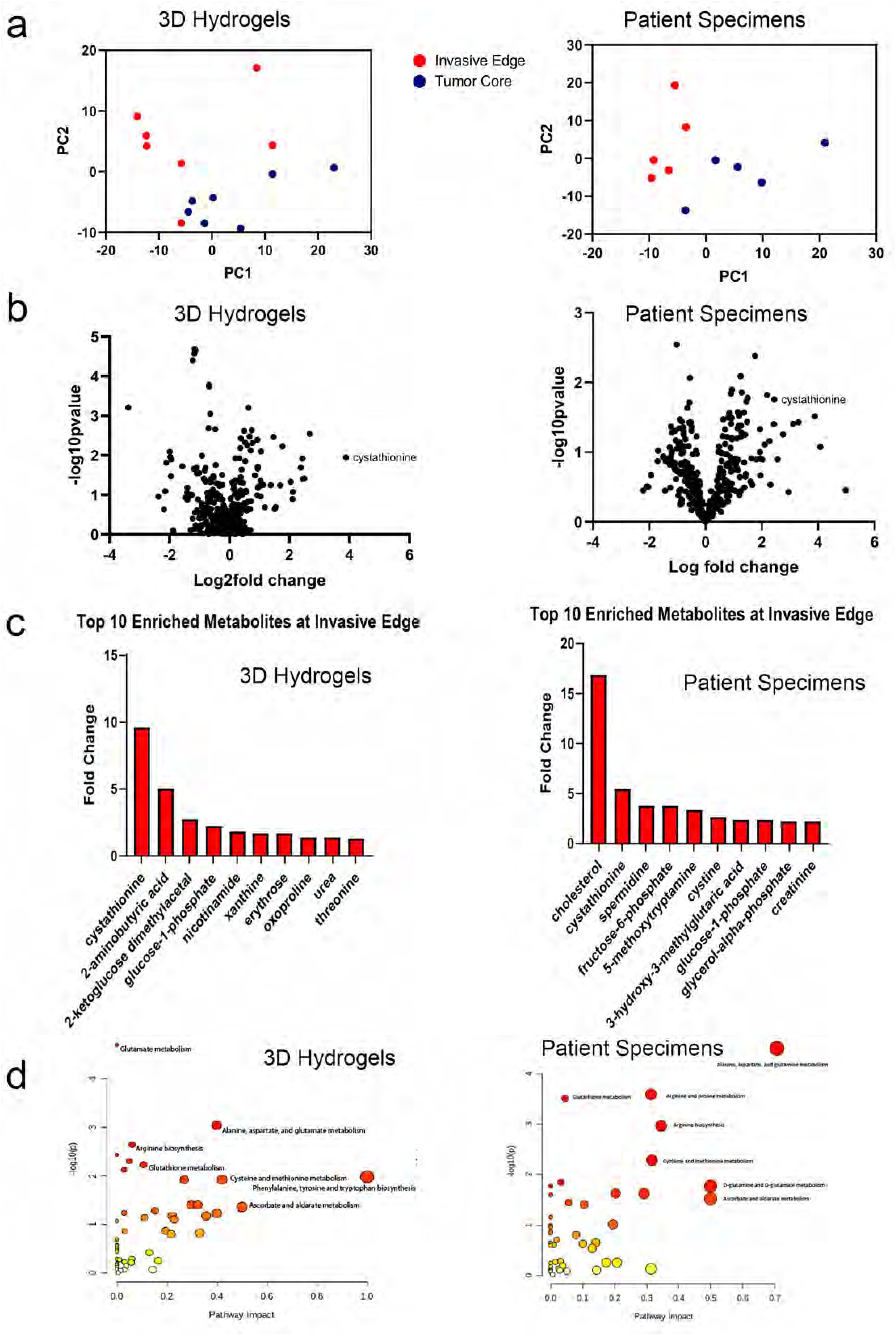
Invasive GBM cells display a distinct metabolic profile in which cystathionine and other oxidative stress metabolites are upregulated. Shown are results from unbiased metabolomic analysis of cells from the invasive front and tumor core of GBM43 cells in 3D hydrogels (left) and site directed biopsies (right) of patient GBMs. (**a**) Principal Component Analysis (PCA) from 3D hydrogels (left; n=7/group) and site directed patient biopsies (right, n=5/group). (**b**) Volcano plots displaying fold-change for metabolites in the invasive front of 3D hydrogels (left) and patient tumors (right) compared with the tumor core. (**c**) Bar graphs displaying 10 most enriched metabolites at the invasive tumor front of 3D hydrogels (left) and patient tumors (right). (**d**) Pathway analysis was performed using MetaboAnalyst to identify pathways upregulated at the invasive tumor front of 3D hydrogels (left) and patient tumors (right). Pathways are plotted according to significance (y-axis) and pathway impact value (x-axis). The node color is based on its p value (darker colors represent more significance) and the node radius is determined based on their pathway impact values (larger circles represent greater pathway enrichment). Most contributing pathways are in the top right corner.

### Lipidomic profiling indicates increased oxidative stress, lipid peroxidation, and apoptotic signaling at the invasive tumor front

To further define metabolic changes associated with GBM invasion, metabolomic analysis was supplemented with high throughput lipidomic analysis of invasive and core GBM cells in 3D hydrogels and patient specimens (**Supplementary Table 2**). Volcano plots (**Fig. 2a**) profiling the 691 identified lipids revealed some overlap, but not perfect agreement, in lipid perturbations between invasive vs. core GBM cells in 3D hydrogels and patient specimens, including elevated glycosylceramides, hexosylceramides, cholesterol esters, and phosphatidylserines, along with reduced triglycerides and acylcarnitines. These trends within lipid groups were revealed in heat maps for 3D hydrogels and patient specimens (**Fig. 2b**). To examine the physiologic role of the differences in lipid abundance between the invasive fraction and tumor core, KEGG metabolic pathway analysis was performed on individual lipids upregulated in invasive GBM cells from both hydrogel models and patient specimens. Pathway enrichment revealed the upregulation of cellular pathways involved in the response to oxidative stress, lipid peroxidation, and anti-apoptotic signaling at the tumor front (**Fig. 2c**). Hexosylceramides and glucosyl ceramides (**Fig. 2d**), which represent modifications of ceramide that cancer cells utilize to prevent ceramide-induced apoptosis occurring during oxidative stress (**Fig. 2e**),^13^ were nearly universally upregulated among the upregulated lipid subgroups in the invasive fractions of hydrogel models and patient GBMs.

**Figure 2.**
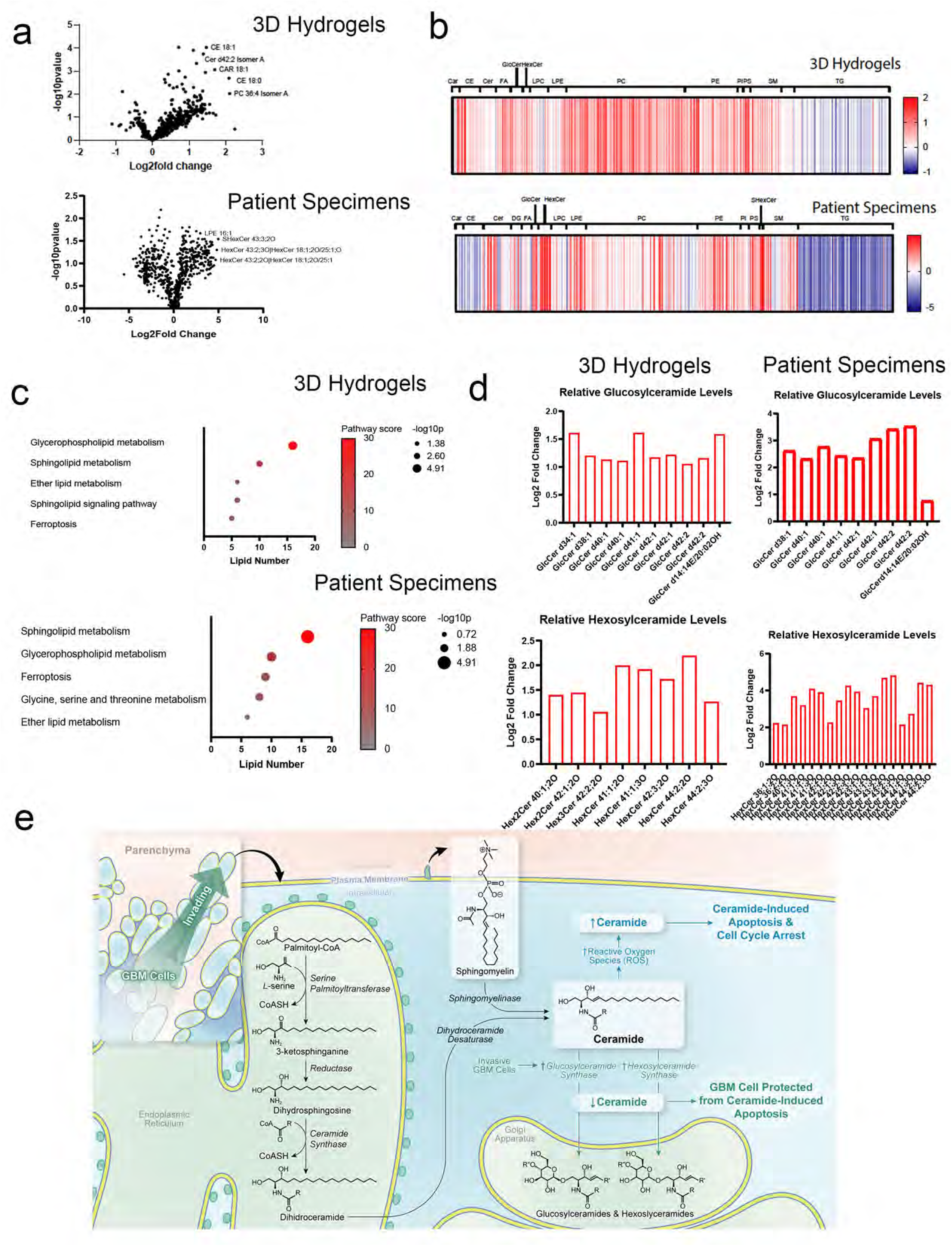
Lipidomic profiling indicates increased oxidative stress, lipid peroxidation, and apoptotic signaling at the invasive GBM front. Shown are results from unbiased lipidomic analysis of cells from the invasive front and tumor core of GBM43 cells in 3D hydrogels and site directed biopsies of patient GBMs. (**a**) Volcano plots displaying relative fold-change for individual lipid abundance at the invasive front of 3D hydrogels (top) and patient specimens (bottom) compared with the tumor core. (**b**) Heat maps displaying relative abundance of lipid species in 3D hydrogels (top) and patient specimens (bottom) organized by lipid classification. (**c**) KEGG pathway enrichment analysis of untargeted lipidomics displaying lipid pathways preferentially upregulated at the invasive tumor front of 3D hydrogels (top) and patient tumors (bottom) using bubble plots. (**d**) Relative fold change of hexosylceramide and glucosylceramide species at the invasive tumor front in 3D hydrogels (left) and patient tumors (right). (**e**) Illustration depicting lipidomic pathways enabling hexosylceramide and glucosylceramide species to protect against apoptosis in invasive GBM cells exposed to oxidative stress.

### Transcriptomic profiling of invasive GBM cells reveals increased expression of genes involved in producing and responding to oxidative stress

To identify gene expression changes associated with the altered levels of hydrophilic metabolites and lipidomes identified by metabolomic and lipidomic analyses, we extracted RNA from invasive and core GBM43 cells from the hydrogel invasion devices. These samples were transcriptomically assessed using the NanoString nCounter panel consisting of a multiplex to analyze the expression of 770 genes across 34 annotated metabolic pathways (**Supplementary Table 3; Extended Data Fig. 2b**). PCA plots of the resulting data revealed that cells in the invasive front clustered together, but apart from cells in the tumor core (**Extended Data Fig. 2c**), indicating a consistent gene expression pattern differentiating cells in the invasive fraction relative to cells in the core fraction. A heatmap (**Extended Data Fig. 2d**) and Volcano plot (**Extended Data Fig. 2e**) revealed the most differentially expressed genes in the invasive fraction relative to the core, which included genes with demonstrated roles in GBM invasion such as *THBS1* (thrombospondin).^14^ We then performed gene set enrichment analysis (GSEA) to determine which processes these genes enriched in GBM43 cells in the invasive fraction relative to the core were involved in. Interestingly, GSEA revealed upregulation of several pathways related to the cellular response to oxidative stress such as the ROS response (**Extended Data Fig. 2f; Fig. 3a**), as well as pathways that generate ROS such as mitochondrial respiration (**Fig. 3a**).

**Figure 3.**
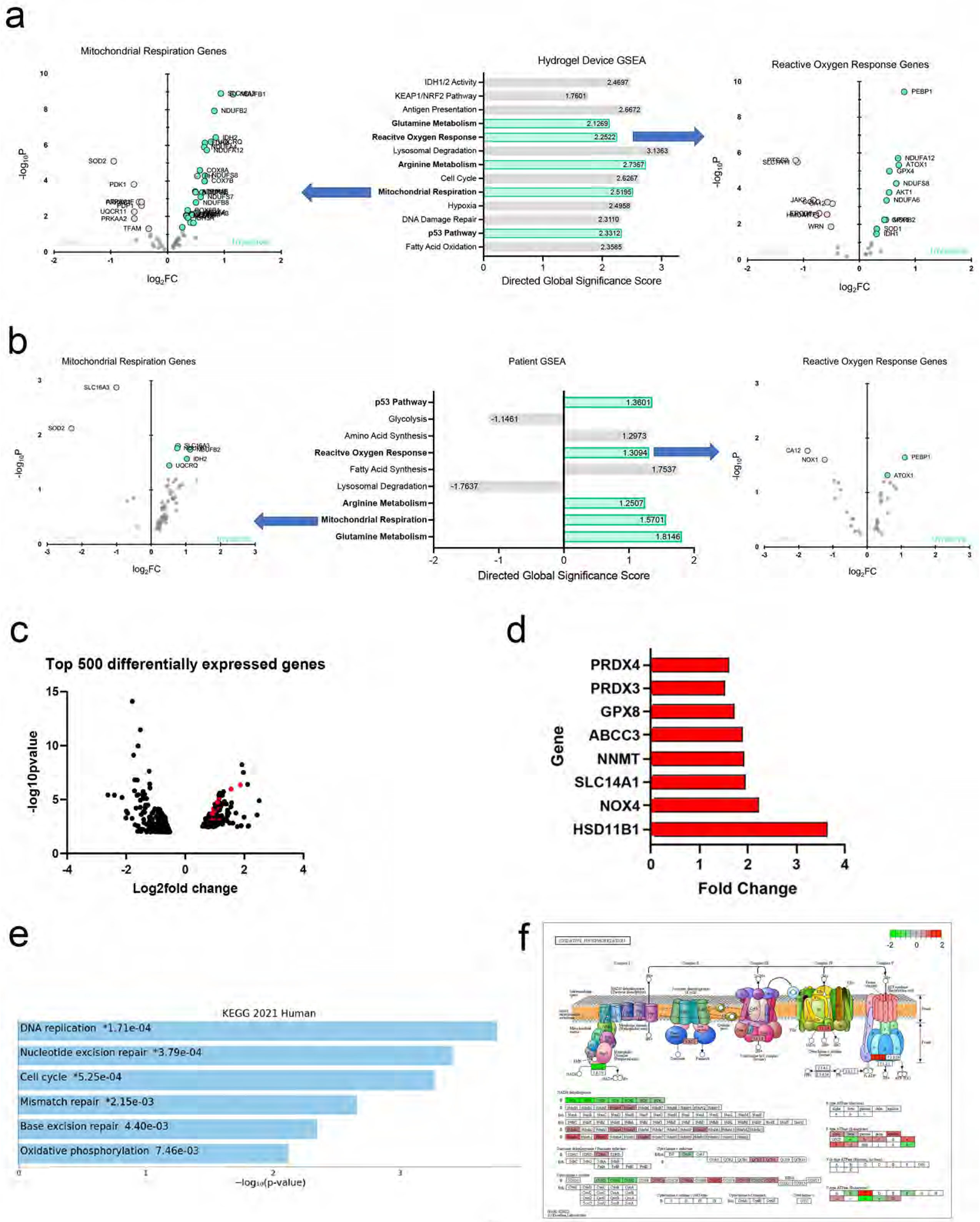
Gene expression profiling demonstrates upregulation of pathways involved in adapting to oxidative stress in invasive GBM cells. (**a-b**) RNA extracted from invasive and core (**a**) GBM43 cells from hydrogel invasion devices or (**b**) site-directed biopsies of patient GBMs were assessed using the NanoString nCounter panel consisting of a multiplex to analyze the expression of 770 genes across 34 annotated metabolic pathways, with GSEA revealing enriched metabolic pathways, including five shared between GBM43 cells in hydrogels and patient specimens (shown in green). Volcano plots (P and FC=probability of significance and fold change invasive vs. core) are shown for genes in two of these pathways - mitochondrial respiration (left) and ROS response genes (right), highlighting genes in the invasive (log_2_FC>0) and core (log_2_FC<0) samples. (**c-f**) We performed bulk RNA sequencing on invasive and core GBM43 cells isolated from our hydrogel invasion devices, revealing that (**c**) of the 250 genes most enriched in the invasive edge relative to the core of hydrogels, 57 (23%) were involved in cellular metabolism; (**d**) among the metabolic genes overrepresented at the invasive edge of the tumors, most were involved in responding to reactive oxidative species (ROS), including *PRDX4*, *PRDX3*, *GPX8*, *ABCC3*, *NNMT*, *SLC14A1*, *NOX4*, and *HSD11B1*; and (e) KEGG pathway analysis implicated pathways involved in the production of and response to ROS rather than pathways related to invasive cellular behavior; and (**f**) gene expression changes overlaid on a schematic of oxidative phosphorylation revealed particular upregulation of genes encoding mitochondrial complex I proteins, a major producer of cellular ROS, in invasive GBM43 cells relative to those in the core.

To determine if these findings were reflective of patient GBMs, the same multiplex platform was then used to analyze metabolic gene expression in RNA extracted from matched specimens taken from the invasive edge and tumor core of patient GBM specimens (n=3) (**Supplementary Table 4**). This analysis yielded a Volcano plot delineating upregulated genes (**Extended Data Fig. 2g**), with GSEA analysis of these upregulated genes revealing that invasive cells in patient GBMs upregulated genes in the amino acid synthesis pathway, including the transsulfuration pathway for cysteine and methionine production whose components were identified in our metabolomic analysis. GSEA analysis of patient GBM specimens from the invasive front also revealed upregulation of five metabolic pathways that were upregulated in invasive GBM43 cells from the 3D hydrogel models, including genes involved in ROS response, mitochondrial respiration, and glutamine metabolism (**Fig. 3b; Extended Data Fig. 2h**). Thus, metabolic transcriptomic analysis revealed increased production of and adaptation to oxidative stress in invasive GBM cells in both hydrogel devices and patient GBM specimens.

To determine how these metabolic gene expression changes fit in with broader transcriptomic changes in invasive GBM cells, we performed bulk RNA sequencing on invasive and core GBM43 cells isolated from our hydrogel invasion devices, revealing the top 250 most enriched genes in cells isolated from the invasive edge versus core of the hydrogel models (**Supplementary Table 5**). Of these 250 genes most enriched in the invasive edge relative to the core of hydrogels, 57 (23%) were involved in cellular metabolism (**Fig. 3c**), underscoring the important role that altered metabolism plays in the broader transcriptomic changes occurring during GBM invasion. Among the metabolic genes overrepresented at the invasive edge of the tumors, the majority were involved in the cellular response to reactive oxidative species (ROS), including *PRDX4*, *PRDX3*, *GPX8*, *ABCC3*, *NNMT*, *SLC14A1*, *NOX4*, and *HSD11B1* (**Fig. 3d**). These transcriptional changes were validated in an additional patient data set when analysis of the Ivy Glioblastoma Atlas Project (Ivy GAP) confirmed elevated expression of 3 of these 8 genes in the invasive front relative to the tumor core (n=10 patients; **Extended Data Fig. 3**). We then analyzed the upregulated pathways from bulk RNA-seq of invasive GBM43 cells in hydrogel devices and found several pathways that could generate ROS, such as oxidative phosphorylation, or that could help cells cope with oxidative stress, such as mismatch repair, nucleotide excision repair, and base excision repair (**Fig. 3e**). A more detailed interrogation of genes involved in oxidative phosphorylation revealed a particular upregulation of genes involved in mitochondrial complex I, which has been particularly implicated in generating ROS^15^ in invasive GBM43 cells in hydrogel devices (**Fig. 3f**).

### Invasive GBM cells exhibited elevated ROS levels

Because multi-omic analysis of invasive GBM cells in hydrogel platforms and patient site directed biopsies reflect a potentially heightened ability to produce and adapt to oxidative stress, we next asked whether invasive GBM cells exhibit higher levels of ROS than core GBM cells. While direct assessment of ROS in tissues is challenging, it is possible to infer ROS presence by measuring biomarkers of oxidative damage which result from the effects of ROS on protein, carbohydrates, nucleic acids and lipids.^16^ We therefore assessed levels of malondialdehyde, a product of fatty acid peroxidation reflecting ROS levels,^17^ and nitrotyrosine, the main product of tyrosine oxidation and a marker of ROS,^18^ in invasive GBM cells in hydrogel devices and in site-directed biopsies of patient GBMs. Immunostaining revealed increased malondialdehyde in the invasive edge compared to the tumor core of 3D hydrogels (P<0.001; **Fig. 4a**) and patient GBMs (P=0.025; **Fig. 4b**). Immunostaining also revealed elevated nitrotyrosine in invasive GBM cells compared to those in the tumor core of hydrogels (P<0.05; **Fig. 4c**), although there was no difference in nitrotyrosine levels between the tumor core and invasive edge of patient specimens (P=0.4; **Fig. 4d**).

**Figure 4.**
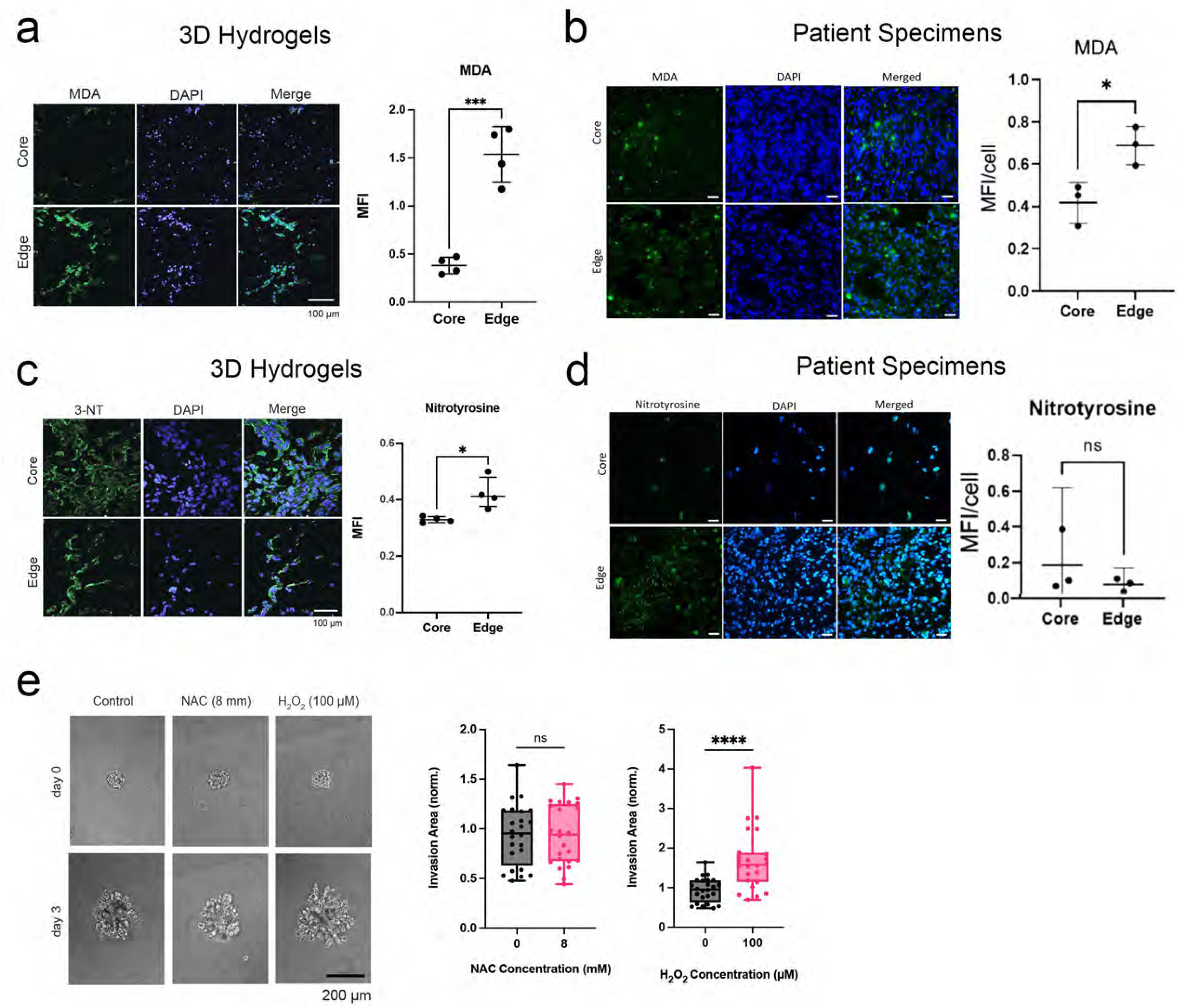
Invasive GBM cells exhibit increased ROS. (**a-b**)Malondialdehyde (MDA) staining of (**a**) hydrogel devices and (**b**) patient specimens revealed increasing MDA in the invasive edge relative to the core of hydrogels (P<0.001; n=4 pairs) and patient specimens (P=0.03; n=3 pairs). (**c-d**) Nitrotyrosine staining of (**c**) 3D hydrogels and (**d**) patient specimens revealed increased staining in the edge relative to the core in the hydrogels (P<0.05; n=4 pairs) but not in the patient specimens (P=0.4; n=3 pairs). (**e**) While hydrogen peroxide increased invasion of GBM43 cells in HA hydrogels (P<0.0001), ROS scavenger N-acetylcysteine did not affect invasion (P=ns) of GBM spheroids in HA hydrogel invasion assays (n=24 spheres, collected across 3 independent experiments).

To determine whether these elevated ROS were capable of promoting invasion rather than merely being a byproduct of the invasive process, we next assessed the impact of ROS manipulation on GBM43 spheroid invasion in hydrogel models (**Extended Data Fig. 4a**), focusing on the three ROS most commonly identified in human cancers (superoxide, hydrogen peroxide, and hydroxyl free radicals).^3^ We found that the ROS hydrogen peroxide increased GBM43 spheroid invasion in the hydrogel models (P<0.001; **Fig. 4e**) at multiple concentrations (**Extended Data Fig. 4b**). Similarly, MnTBAP, a metalloporphyrin superoxide dismutase (SOD) mimetic which lowers superoxide (P<0.001; **Extended Data Fig. 4c**) by converting it to peroxide also increased GBM43 spheroid invasion in HA hydrogels (P<0.001; **Extended Data Figs. 4d-e**). In contrast, N-acetylcysteine did not affect invasion (P>0.05; **Fig. 4e**) at multiple concentrations (**Extended Data Fig. 4b**), but also did not affect superoxide levels in GBM43 cells (**Extended Data Fig. 4f**). This suggests that, amongst the ROS most commonly identified in human cancers, hydrogen peroxide promoted GBM invasion in hydrogel platforms.

### CRISPR screen of metabolic genes links transsulfuration pathway to invasion

To determine which upregulated metabolic pathways identified by our multi-omics analysis enable GBM cell invasion, we performed a CRISPR-Cas9 knockout (KO) screen on a library of 29790 sgRNAs targeting 2981 metabolic genes^19^ to identify metabolic genes crucial to GBM invasion. GBM43 cells expressing Cas9 and the sgRNA library were seeded in 3D hydrogel invasion devices (n=6) and cultured for 28 days. Afterwards, devices were disassembled and hydrogels were microdissected to isolate invasive and core cells for subsequent DNA sequencing (**Extended Data Fig. 5a**). sgRNAs enriched in the core relative to the invasive fraction (indicating genes whose knockdown disrupted invasion) and in the invasive fraction compared to the core (indicating genes whose knockdown enabled invasion) were scored based on their abundance compared to non-targeting sgRNAs in the library (**Supplementary Tables 6-8; Extended Data Fig. 5b**) We chose five genes (*SMS*, *NDUFS8*, *SMPD1*, *CTH*, and *COMDT*) based on the enrichment of sgRNAs targeting them in the core (**Fig. 5a; Supplementary Table 8**) and their overlap with our transcriptomic, metabolomic, and lipidomic datasets. We then performed single gene knockdowns of these five genes using CRISPRi to test the effect of gene silencing on tumor spheroid invasion (**Extended Data Fig. 5c**). All knockdown cell lines were tested in HA hydrogel spheroid invasion assays to assess invasive capacity and compared to control GBM43 cells expressing dCas9. All five knockdown cell lines exhibited decreased spheroid invasion in HA hydrogels compared to control spheroids (P<0.001; **Fig. 5b**).

**Figure 5.**
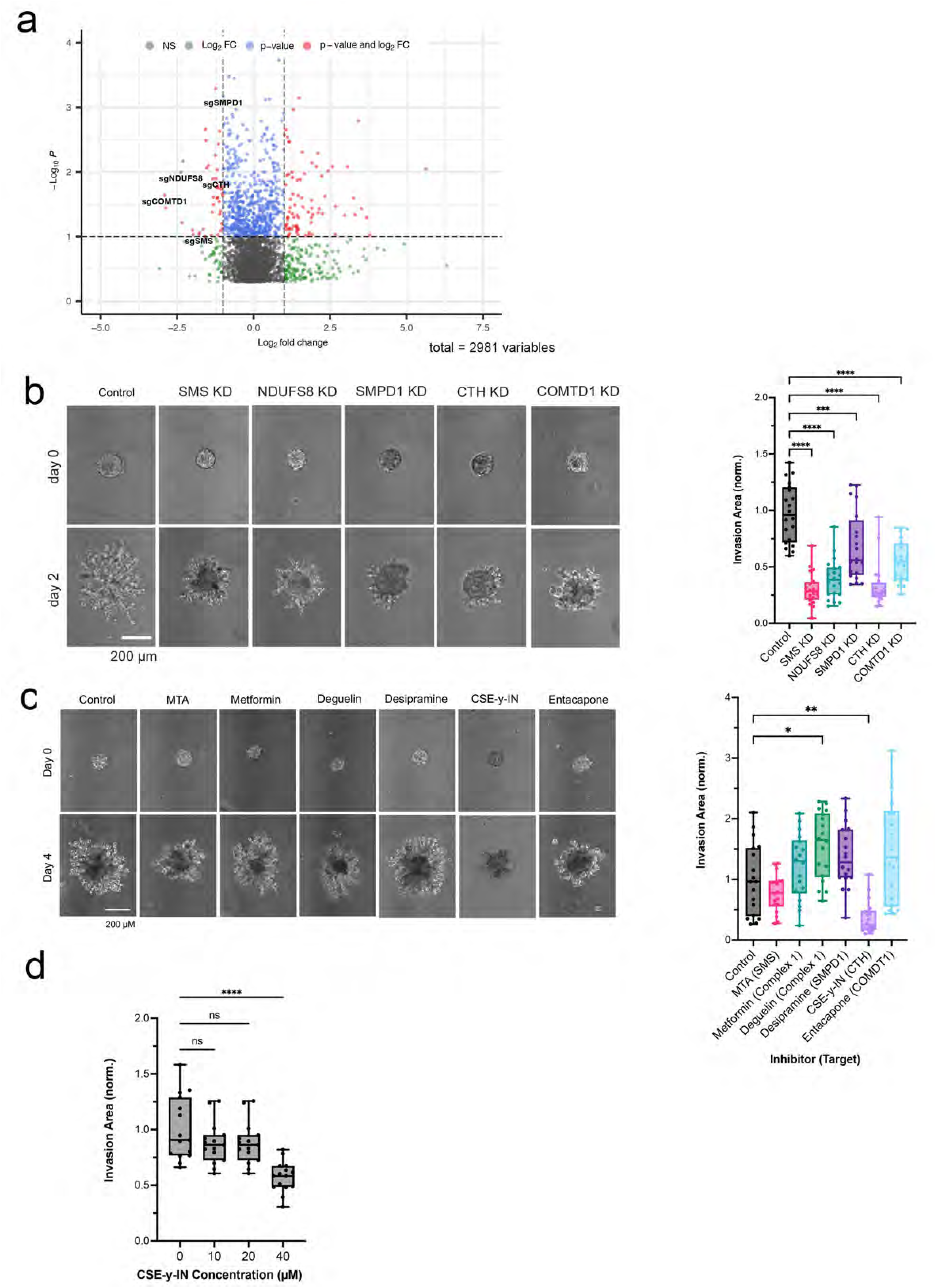
A focused metabolic CRISPR screen to identify metabolic genes essential to GBM invasion reveals that ROS response genes including CTH are necessary for tumor cell invasion. (**a**) Shown is a volcano plot displaying the enrichment of sgRNAs for metabolic genes in the core (log_2_fold change < 0) and invasive front (log_2_fold change > 0) of GBM 3D invasion devices, with labeling of the five genes (*COMTD1*, *SMS*, *CTH*, *SMPD1*, and *NDUFS8*) selected for further evaluation. (**b**) Quantification and representative images of spheroid invasion assays of five knockdown GBM43 cell lines selected from CRISPR screen hits compared with control cells expressing dCas9 (n=20 spheres, collected across 3 independent experiments). (**c**) Spheroid invasion assays of GBM43 cells treated with inhibitors of the five metabolic enzymes encoded by the genes chosen for further evaluation from the CRISPR screen (n=18 spheres, collected across 3 independent experiments). (**d**) CSE-γ-IN, an inhibitor of CTH, slowed GBM43 tumorsphere invasion at 40 μM (P<0.0001; n=15 spheres, collected across 3 independent experiments).

We then assessed the effect of pharmacologically inhibiting the products of these five genes and their associated pathways on GBM43 spheroid invasion assays. Only one inhibitor, cystathionine-y-lyase-IN-1 (CSE-γ-IN), slowed invasion (P<0.01; **Fig. 5c**). CSE-γ -IN is a small molecule inhibitor of cystathionine gamma lyase (CTH/CSE), an enzyme catalyzing cystathionine breakdown into cysteine in the last step of the transsulfuration pathway,

To determine whether these inhibitory effects on invasion were attributable to an effect on cell survival, we also assessed the concentration window for which CSE-y-IN inhibited GBM spheroid invasion without cytotoxicity and found that the invasion inhibitory effect of CSE-γ-IN on GBM43 spheroids began at 40 μM (P<0.0001; **Fig. 5d**) with 40 μM CSE-γ-IN also inhibiting U251 spheroid invasion (P<0.001; **Extended Data Fig. 5d**). Concentrations of CSE-γ-IN above 100 μM began to affect the viability of GBM43 and U251 cells (**Extended Data Fig. 5e**).

The ability of CTH to play a functional role in invading GBM cells was further supported by our finding from Ivy GAP analysis that pyridoxal kinase (PDXK), the enzyme that converts pyridoxine and other vitamin B6 precursors into pyridoxal-5′-phosphate (PLP), the bioactive form of CTH cofactor vitamin B6,^20^ was enriched at the leading edge of the tumor relative to the core (P<0.001; **Extended Data Fig. 5f**).

### Transsulfuration pathway inhibition slows GBM invasion

Because CTH was the only metabolic gene emerging from our CRISPR screen whose pharmacologic targeting inhibited invasion in spheroid invasion assays (**Fig. 5c**) and because cystathionine, a precursor to cysteine in glutathione synthesis in the transsulfuration pathway, was enriched in the invasive fraction of both patient derived tumor biopsies and 3D hydrogels (**Fig. 1c**), we focused further investigation on the specific role of CTH in GBM invasion. We first expanded upon the effects of CTH knockdown on invasion by demonstrating that CTH knockdown slowed long-term GBM43 invasion in 3D hydrogel devices (28 day culture period; **Fig. 6a; Extended Data Fig. 6a**). CTH knockdown led to higher ROS levels in normoxia but not in hypoxia, as detected by the CellROX reagent which measures hydroxyl radical and superoxide anion,^21^ in cultured GBM43 cells (P=0.01-0.03; **Extended Data Fig. 6b**). Growth in low concentrations of cysteine (200 µM) amplified ROS levels in CTH knockdown and control cells while preserving the elevated ROS seen in CTH knockdown (P<0.001; **Extended Data Fig. 6c**). CTH knockdown did not alter levels of superoxide in cultured GBM43 cells in normal (400 μM) and low (200 μM) cysteine concentrations as measured by the MitoSOX^TM^ probe (P=0.8 at 400 μM cysteine; P=0.2 at 200 μM cysteine; **Extended Data Fig. 6d**), suggesting that hydroxyl radical accumulates in cells deprived of cysteine due to *CTH* knockdown. These results are consistent with cell-free chemistry studies implicating cysteine disulfides in the antioxidant response to hydroxyl radical attack.^22^ Interestingly, levels of ROS assessed by the CellROX^TM^ reagent did not differ between the core and invasive fractions of GBM43 control and CTH knockdown cells in 3D hydrogel devices (**Extended Data Figs. 6e-f**), possibly reflecting greater hypoxia in the devices.

**Figure 6.**
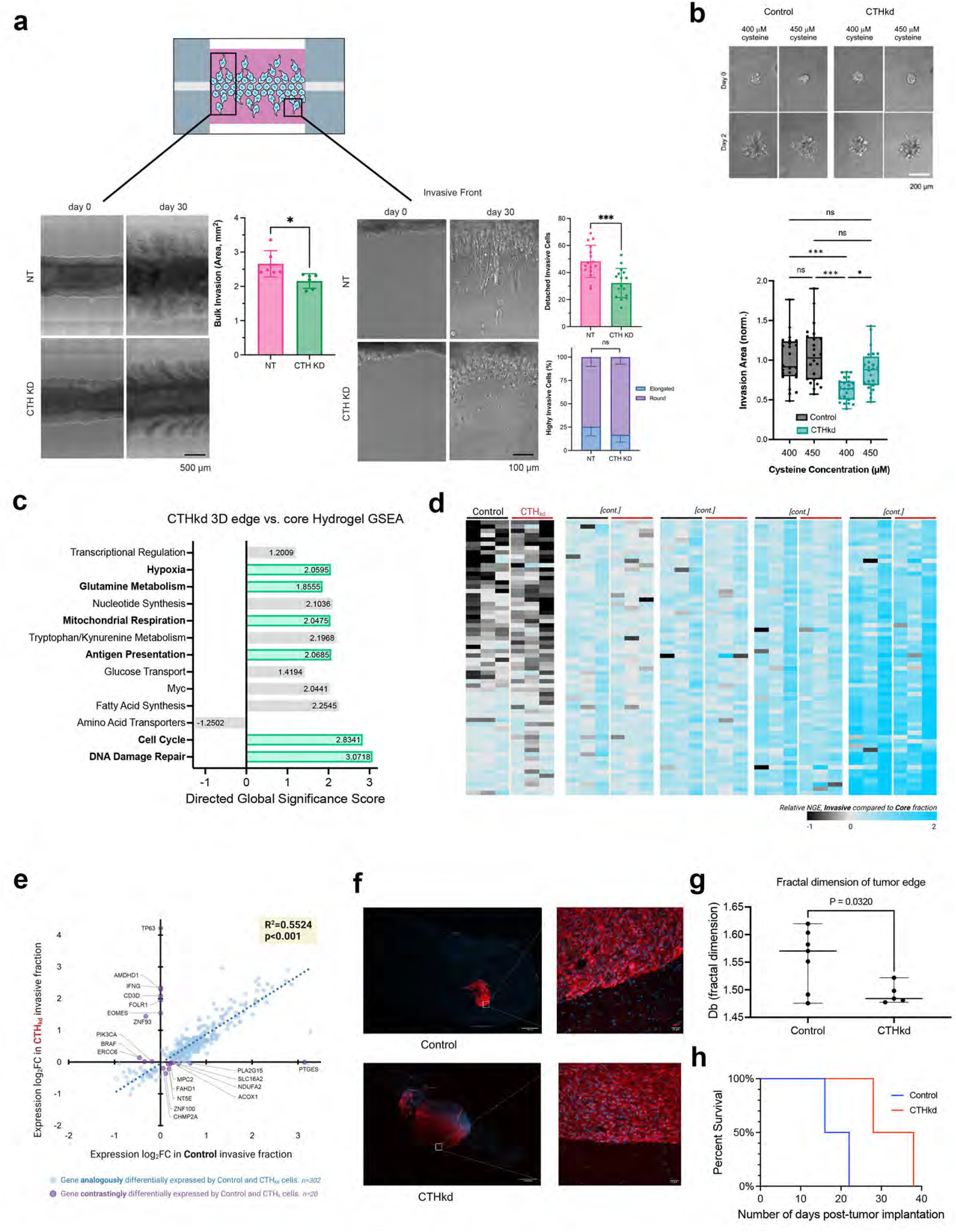
Targeting of cystathionine gamma lyase (CTH) inhibits GBM invasion. (**a**) GBM43 cells with CRISPRi targeting of CTH were validated to be less invasive in 3D hydrogel invasion devices as determined by bulk invasive area (left; P<0.05; n=6 regions of interest, collected across 3 devices) and number of detached invasive cells (right; P<0.001), with morphology of invasive cells not affected by CTH knockdown (right; P>0.05; n=16 regions of interest, collected across 3 devices). (**b**) Spheroid invasion assays revealed that additional cysteine supplementation (increasing from 400 μM in normal media to 450 μM) reversed the slowed invasion caused by CTH knockdown (n=24 spheres, collected across 3 independent experiments). (**c-e**) GBM43 cells with CRISPRi knockdown of CTH were seeded into invasion devices, after which cells isolated from core and invasive fractions were transcriptomically assessed using the NanoString nCounter platform and the 770 metabolic gene multiplex, yielding (**c**) a GSEA in which 6 of the 13 upregulated pathways were shared with control cells invading hydrogel devices (shown in green); (**d**) a heatmap depicting normalized gene expression (NGE) of cells in the invasive fraction relative to core fraction from same hydrogel for CTHkd (red bars) and control GBM43 cells (black bars) (n=3 per group), with uniform gene expression changes across control GBM43 vs. CTHkd cells suggesting similar transcriptional profile in this platform amongst invasive GBM cells regardless of CTH expression; and (**e**) scatter plot depicting the fold-change in gene expression for the same genes, in invasive compared to core fractions for GBM43 control cells (x-axis) and GBM43 CTHkd (y-axis). The high correlation between fold-change values in invasive control GBM43 vs. CTHkd cells (P<0.001) illustrates that transcriptional patterns related to metabolism change during invasion similarly regardless of CTH expression. Purple dots indicate genes with discordant expression changes in control GBM43 vs. CTHkd cells, which are scant (20 of 322 total genes=6.2%). (**f-h**) Intracranial PDXs derived from GBM43 cells expressing mCherry along with dCas9 or dCas9 with sgRNAs targeting CTH (**f-g**) were less invasive with CTH knockdown (P=0.03; n=5-7/group) based on fractal analysis of images of tumors and their surrounding brain, which yields the fractal dimension, a numeric description of invasive tumor growth pattern as a continuous number between 1 and 2, with higher numbers representing greater invasiveness (10x and 20x magnification with 1000 μm and 25 μm scale bars); and (**h**) exhibited prolonged survival with CTH knockdown (P=0.008; n=4/group).

### The transsulfuration pathway is necessary for GBM invasion because of its role in *de novo* cysteine synthesis

Because the transsulfuration pathway is the primary route for cysteine and glutathione biosynthesis, we next investigated whether GBM cells lacking CTH are less invasive due to a limited supply of either of these two major antioxidants produced downstream of CTH. We performed a spheroid invasion assay in the presence of 50 µM additional cysteine to see if exogenous cysteine can reverse the decreased invasion of CTH knockdown cells. In standard culture media containing 400 µM cysteine, GBM43 spheroids with CTH knockdown were less invasive than control spheroids; however, increasing total cysteine to 450 µM “rescued” the ability of GBM43 spheroids with CTH knockdown to invade to a level similar to that of control GBM43 spheroids (P=0.01; **Fig. 6b**). To test the possibility that cysteine promotes invasion by serving as a precursor to glutathione, we also performed invasion assays with control and CTH knockdown cells with glutathione supplementation. Surprisingly, glutathione supplementation did not rescue the invasive ability of CTH knockdown cells (P>0.05; **Extended Data Figs. 7a-b**). In fact, GBM43 CTH knockdown cells had higher levels of glutathione than control GBM 43 cells (P=0.01-0.03; **Extended Data Fig. 7c**), suggesting that alternative pathways of glutathione production were circumventing any impact of CTH knockdown on glutathione levels and that cysteine was driving invasion through glutathione-independent pathways.

Recent studies have defined ferroptosis as a form of cell death characterized by the accumulation of the types of lipid peroxides^23^ that we found invasive GBM cells to be protected against (**Fig. 2e**). We therefore investigated whether cysteine supplementation or CTH targeting affected the sensitivity of GBM cells to erastin, a small molecule that induces ferroptotic cell death. We found that additional cysteine supplementation protects GBM43 cells from erastin-induced cell death (P<0.001) and inhibiting the transsulfuration pathway with CSE-γ-IN (P<0.05) makes the cells more sensitive to erastin (P<0.05; **Extended Data Figs. 7d-e**).

### Upregulation of other transsulfuration enzymes in GBM cells invading despite CTH knockdown reveals the importance of cysteine for GBM invasion

While CTH knockdown considerably slowed GBM43 invasion through 3D hydrogels in spheroid invasion assays and long-term invasion devices (**Fig. 5b**), a population of GBM43 CTH knockdown cells remained moderately invasive in our assays. We therefore investigated whether invasive CTH knockdown GBM43 cells relied on similar metabolic genes and pathways as invasive control GBM43 cells or if they utilized compensatory pathways to invade. First, we performed Ki-67 staining of the core and invasive fractions of GBM43 control and CTH knockdown cells cultured in the invasion devices. There were no differences in the percentage of Ki-67 positive cells between the control and CTH knockdown cells in either the core or invasive fractions of the devices (**Extended Data Fig. 8a**), confirming that differences in invasion between the cell lines did not reflect proliferative differences. Then, cells isolated from core and invasive fractions were transcriptomically assessed using the NanoString nCounter platform and the same 770 metabolic gene multiplex described above (**Supplementary Tables 9-10; Extended Data Fig. 8b**). PCA plots revealed that cells in the invasive front clustered together, but apart from cells in the core (**Extended Data Fig. 8c**). A Volcano plot (**Extended Data Fig. 8d**) and heatmap (**Extended Data Fig. 8e**) were created to identify enriched metabolic genes (**Extended Data Fig. 8f**) in the invasive fractions relative to the core fractions of the hydrogels, with GSEA revealing that genes involved in fatty acid synthesis, glucose transport, tryptophan metabolism, mitochondrial respiration, glutamine metabolism, and hypoxia response were among the gene sets most enriched in the invasive cells (**Fig. 6c**).

We then compared these upregulated metabolic genes in invasive CTH knockdown GBM43 cells to those upregulated in invasive control GBM43 cells. A heatmap revealing DEGs (differentially expressed genes) across invasive samples from CTH knockdown versus control GBM43 cells revealed an unchanged general pattern of metabolic gene expression between CTH knockdown versus control GBM43 cells (**Fig. 6d**). Similarly, a scatter plot comparing the fold-change in gene expression for individual genes in invasive compared to core fractions of both cell lines (CTH knockdown and control) revealed a high correlation between fold-change values in CTH knockdown versus control samples (P<0.001), with only 6.2% (20/322) of the genes having discordant expression changes in CTH knockdown versus control cells in the invasive fractions relative to core fractions (**Fig. 6e**). This finding was also corroborated in a Volcano plot demonstrating negligible differences in upregulated genes within the 770 metabolic gene multiplex in the invasive fraction compared to the core in CTH knockdown vs. control GBM43 cells (**Extended Data Fig. 8g**). Together, these findings revealed that transcriptional patterns related to metabolism change similarly regardless of CTH expression.

Having demonstrated no differences in expression of the 770 metabolic genes in the multiplex between invasive CTH knockdown vs. control GBM43 cells, we then expanded this transcriptomic comparison using bulk RNA-seq and found that invasive CTH KD cells exhibited upregulation of another enzyme in the transsulfuration pathway, CBS (cystathionine beta-synthase) (**Extended Data Fig. 8f; Supplementary Tables 11-13**). Because CBS catalyzes the first step of transsulfuration by condensing serine with homocysteine to generate cystathionine upstream of CTH generating cysteine from cystathionine, our finding of CBS upregulation in invasive CTH KD cells underscored the essential nature of cysteine for GBM invasion.

### CTH knockdown slows GBM invasiveness *in vivo*

Finally, we analyzed the invasiveness of intracranially implanted GBM43 cells with or without CTH knockdown (**Extended Data Fig. 9a**). Invasiveness *in vivo* was assessed by fractal analysis of images of tumors and their surrounding brain, which yields the fractal dimension, a numeric description of invasive tumor growth pattern as a continuous number between 1 and 2, with higher numbers representing greater invasiveness. Application of this method to intracranial GBM43 tumors with or without CTH knockdown revealed that CTH knockdown reduced PDX invasiveness (P=0.03; **Figures 6f-g**), with associated increase in survival (P=0.008; **Figure 6h**) despite creating larger tumors (P=0.02; **Extended Data Fig. 9b**). This finding suggests that the improved survival in CTH knockdown PDXs was due to reduced invasiveness as evidenced by examples of tumor metastasizing to the brainstem in mice with GBM43 cells lacking CTH knockdown (**Extended Data Fig. 9c**).

## DISCUSSION

While the hallmark of GBM and a defining contributor to its poor prognosis is invasion into the surrounding white matter, studies to date have focused more on mechanisms driving this invasion rather than the requirements needed to sustain it. To close this knowledge gap, we developed a bioengineered 3D hydrogel invasion platform for high throughput screening of invasion mediators. The novel spatially dissectable nature of our hydrogel-based invasion devices allowed us to perform a multi-omic analysis of tumor cells in the invasive front versus non-invasive core from individual invasion assays. We then benchmarked these findings from our 3D hydrogel models against site-directed biopsies from the core versus invasive edge of patient GBMs. This study represents one of the closest integrations to date of 3D biomaterial models and patient data and illustrates the value of reductionist paradigms for identifying biomarkers and mechanisms of cancer invasion.

By focusing on the previously understudied role of metabolic reprogramming in GBM cellular invasion, we produced several novel insights into this process. First, we found that the invasive front of GBM produces elevated ROS and that invasive tumor cells exhibit metabolomic, lipidomic, and transcriptomic profiles reflective of their exposure to this oxidative stress. Second, we found that hydrogen peroxide is the specific ROS promoting invasion in GBM cells, building upon prior reports linking ROS to the invasiveness of other cancer types.^24^ Third, our unbiased metabolic CRISPR knockout screen identified candidate genes and pathways regulating invasion, with five genes upregulated in invasive GBM cells validated by clonal KD cell lines (*SMS*, *NDUFS8*, *SMPD1*, *CTH*, and *COMDT)*. In particular, we found that inhibiting CTH, an enzyme in the transsulfuration pathway which produces the non-essential amino acid cysteine, impairs the ability of GBM cells to invade through HA hydrogels and that this impaired invasion can be “rescued” through cysteine supplementation. These results suggest that mitigation of oxidative stress is crucial for GBM invasion and that this process can be disrupted by targeting the adaptive response to oxidative stress, including the transsulfuration pathway. Our finding that cysteine deficiency drives the diminished invasiveness caused by CTH knockdown cells builds upon prior studies in which cysteine depletion induces ferroptosis in other cancers^25^ by suggesting that cysteine depletion slows GBM invasion by impairing the ability of invasive GBM cells to cope with oxidative stress and evade ferroptosis.

Several of our findings implicated the transsulfuration pathway, which synthesizes the non-essential amino acid cysteine via the intermediate cystathionine, as being critical for GBM invasion. First, cystathionine, the central metabolite in the transulfuration pathway, was among the two most enriched metabolites in the invasive edge of 3D hydrogels and GBM patient specimens, with 2-aminobutyric acid, a byproduct of cystathionine conversion to cysteine,^26^ the second most enriched metabolite in the invasive edge of 3D hydrogels. Second, CTH, a central enzyme in the transsulfuration pathway was identified by a metabolic CRISPR screen to be crucial for invasion of GBM tumor cells in culture. Knockdown of CTH confirmed its importance in the invasive process for GBM cells and suppressed tumor cell invasion to a greater extent than the other validated hits from the CRISPR screen.

Interestingly, in healthy brain, the transsulfuration pathway has been confirmed to be intact but inefficient at later steps, with cystathionine present at higher levels in the brain compared to other organs.^27^ This inefficiency is then exacerbated in neurodegenerative conditions such as Parkinson’s and Alzheimer’s diseases.^27^ Our studies suggest that invasive GBM cells upregulate and rely on the transsulfuration pathway beyond the level seen in healthy brain to generate cysteine, enabling them to cope with the oxidative stress associated with the invasive process identified by us in patient GBMs and identified by others in prostate cancer.^24^

Our analysis suggests that the importance of the transsulfuration pathway and CTH in particular in GBM invasion likely arises from its production of the antioxidant cysteine, rather than the antioxidant glutathione which is produced downstream of cysteine and has alternative pathways for its production.^28^ Prior studies have suggested that growing cancer cells have an increased demand for cysteine that exceeds the amino acid’s availability in the adjacent microenvironment.^29–32^ This causes the transsulfuration pathway to be activated to meet the metabolic requirements of the cancer cells.^31,32^ It is logical to hypothesize that a similar phenomenon occurs in GBM cells invading adjacent tissue, where an increased demand for cysteine likely arises from the ROS-induced oxidative stress we identified at the invasive GBM front, with the transsulfuration pathway providing this cysteine due to insufficient cysteine available in the invaded white matter for uptake through membrane transporters.^26,31,32^

The ROS at the invasive GBM edge likely derive from a combination of intrinsic and extrinsic sources.^33,34^ Intrinsically, the biochemical cellular processes which drive invasion in GBM cells could generate a significant amount of ROS, a possibility supported by our finding of increased expression of the components of mitochondrial oxidative phosphorylation most associated with ROS production at the invasive front. Thus, for GBM cells to continue invasion within the brain, the ROS generated by invasion would have to be detoxified.^35,36^ Extrinsically, it is also possible that the abundance of ROS at the invasive tumor edge is simply a product of greater oxygen at the tumor edge than the relatively hypoxic core.^34,37^

Having shown that invasive GBM cells produce oxidative stress and must cope with this stress as part of the invasive process, we then investigated whether these ROS directly promote invasion Prior reports have highlighted the importance of ROS in the metastatic cascade, as well as in numerous other cellular processes across cancer types including tumor proliferation, suppression of apoptotic death, and angiogenesis.^33–35^ The extent to which ROS play a role in tumor cell invasion, however, has not been investigated in depth.^36^ Our studies address this knowledge gap by demonstrating that exposure to hydrogen peroxide, one of three predominant ROS found in cancers like GBM,^38^ increased invasion in 3D hydrogels. This finding has translational implications that should be accounted for when considering therapeutic strategies such as metalloporphyrin SOD mimetics which are in clinical trials (NCT02655601) for GBM, since we found that the ability of these agents to convert superoxide to peroxide promoted GBM invasion.

Further work will also be needed to determine how peroxide drives GBM cell invasion.^35,36^ Previous studies have implicated transcription-independent mechanisms promoting degradation of proteins suppressing invasion via the ubiquitin-proteasome pathway as a mechanism of ROS-promoted lung cancer invasion.^39^ Furthermore, physiologic processes regulated by ROS in other tumor cell types are often dependent on ROS levels, where moderate ROS levels induce a cascade of cellular/extra-cellular signals which promote tumor growth and survival, and high levels of ROS ultimately induce tumor cell apoptosis.^35,37,40^ Indeed, we found a similar dose-response relationship between peroxide concentration and GBM invasion (**Extended Data Fig. 4b**).

Overall, this work demonstrates that invading GBM cells undergo metabolic adaptations to the oxidative stress that invasion elicits, with the transsulfuration pathway enzyme CTH a targetable driver of this invasive metabolic phenotype. We found that the non-essential amino acid cysteine is crucial to the ability of GBM cells to manage the oxidative stress associated with invasion, and the transsulfuration pathway is crucial to cysteine production when its demand exceeds its supply in the tumor microenvironment.

Future studies are needed to investigate the therapeutic potential of targeting GBM invasion via its metabolic dependence on the transsulfuration pathway. Notably, we found upregulation of CBS, the enzyme one step upstream of CTH, in GBM cells invading despite CTH knockdown, suggesting that targeting the transsulfuration pathway at multiple steps may be needed to guard against invasive escape from the targeting of a single step. We also found that, while CTH knockdown improved survival of intracranial tumor-bearing mice *in vivo*, the resulting tumors reached a larger size, consistent with a “go or grow” hypothesis that has been postulated for GBM,^41^ in which invasive and proliferative programs toggle back and forth. This finding would suggest that the benefit of targeting invasion via the transsulfuration pathway may be enhanced if this approach is supplemented with traditional cytotoxic chemotherapy targeting proliferating cells.

## METHODS

### Cell Culture

U-251 MG (University of California, Berkeley Tissue Culture Facility, which sources from ATCC) GBM cells were cultured in DMEM (Thermo Fisher Scientific) supplemented with 10% (vol/vol) fetal bovine serum (Corning, MT 35-010-CV), 1% (vol/vol) penicillin-streptomycin (Thermo Fisher Scientific), 1% (vol/vol) MEM non-essential amino acids (Thermo Fisher Scientific), and 1% (vol/vol) sodium pyruvate (Thermo Fisher Scientific). GBM43 cells (Mayo clinic) were cultured in DMEM (Thermo Fisher Scientific) supplemented with 10% (vol/vol) fetal bovine serum (Corning, MT 35-010-CV), 1% (vol/vol) penicillin-streptomycin (Thermo Fisher Scientific) and 1% (vol/vol) Glutamax (Thermo Fisher Scientific, 35-050-061). Cells were harvested using 0.25% Trypsin-EDTA (Thermo Fisher Scientific) and passaged less than 30 times. All cells were screened on a bimonthly basis for mycoplasma and validated every six months by Short Tandem Repeat (STR) analysis at the University of California Cell Culture Facility.

### 3D Hydrogel Platforms

#### Me-HA Synthesis

Hyaluronic Acid (HA) hydrogels were synthesized as previously described.^42^ Briefly, methacrylic anhydride (Sigma-Aldrich, 94%) was used to functionalize sodium hyaluronate (Lifecore Biomedical, Research Grade, 66 kDa – 99 kDa) with methacrylate groups (Me-HA). The extent of methacrylation per disaccharide was quantified by ^1^H nuclear magnetic resonance spectroscopy (NMR) and found to be ∼85% for materials used in this study. To add integrin-adhesive functionality, Me-HA was conjugated via Michael Addition with the cysteine-containing RGD peptide Ac-GCGYGRGDSPG-NH2 (Anaspec) at a concentration of 0.5 mmol/l.

#### HA Hydrogel Rheological Characterization

Hydrogel stiffness was characterized by shear rheology via a Physica MCR 301 rheometer (Anton Paar) with an 8-mm parallel plate geometry for γ = 0.5% and f = 1 Hz. Frequency was controlled to be between 50 and 1 Hz for the frequency sweep at a constant strain (γ=0.5%), and the modulus saturation curve with time was obtained under oscillation with constant strain (γ=0.5%) and frequency (f = 1 Hz). The temperature of the gel solution was controlled (T = 37°C) with a Peltier element (Anton Paar) and the sample remained humidified throughout the experiment.

#### Tumorsphere Invasion Assays

Tumorspheres were fabricated using Aggrewell Microwell Plates (Stemcell Technologies). Briefly, 1.2×10^5^ cells were seeded into a single well of the Aggrewell plate to form spheroids consisting of 100 cells. After a 48-hour incubation, spheroids were resuspended in phenol red-free serum-free Dulbecco’s Modified Eagle’s Medium (DMEM, Thermo Fisher Sci, 21-063-029) at a density of 1.5 spheroids/μL and used as solvent for HA hydrogel crosslinking. To form hydrogels, 6 wt. % Me-HA was crosslinked in phenol red-free serum-free Dulbecco’s Modified Eagle’s Medium (DMEM, Thermo Fisher Sci, 21-063-029) with a protease-cleavable peptide (KKCG-GPQGIWGQ-GCKK, Genscript). HA-RGD gels were crosslinked with peptide crosslinkers at varying ratios to yield hydrogels with a shear modulus ∼300 Pa and a final 1.5 wt. % Me-HA (Supp. Fig. 1). For all experiments, unless otherwise mentioned, a concentration of 3.405 mM peptide crosslinker was selected to yield a shear modulus of 300 Pa. After 1 hour crosslinking in a humidified 37°C chamber, complete cell culture media was added to the hydrogels. For all experiments unless otherwise noted, cell culture media was replenished every 2 days.

#### Invasion Devices

To fabricate the invasion device, the device base, lid and spacers were laser-cut out of 1.5 mm thick CLAREX° acrylic glass (Astra Products, Inc.). The pieces were assembled and fastened with epoxy, UV-treated for 10 min, and stored in cold room prior to use. On the day of experiment, devices were brought to room temperature and a 22G X 1.5” bevel needle (BD Precision Glide) was inserted into the device to serve as a channel mold. HA hydrogel solution was casted around the wire and incubated for one hour in a humidified 37°C chamber. After crosslinking, devices with hydrogels and needles were submerged in cell culture media for at least 10 min, before removal of the needle which left an open channel. Afterwards, four million cells were seeded into the open channel and the channel ends were plugged with vacuum grease. Unless otherwise stated, invasion devices were cultured for 28 days and media was replenished every 3 days, with 28 days chosen as an ideal long-term culture based on previous experiments revealing it to be the time frame at which control GBM43 cells fully invade through the hydrogel device.

### Invasion Quantification

For invasion analysis of spheroids embedded in HA hydrogels, spheroids were imaged every two days using Eclipse TE2000 Nikon Microscope with a Plan Fluor Ph1×10 objective. Images were acquired using NIS-Elements Software. For each spheroid, invasion was calculated using the following equation: [(A_f_ – A_i_)/A_i_) where A_f_ = final spheroid area and A_i_ = initial spheroid area. Spheroid area was measured in ImageJ and invasion was normalized to control spheroids.

For invasion analysis of cells growing in devices, cells in devices were imaged every seven days using Eclipse TE2000 Nikon Microscope with a Plan Fluor Ph1 ×10 objective. Images were acquired and stitched using NIS-Elements Software. For each device, total cell reservoir area was outlined in ImageJ at each timepoint, and invasion was calculated using the same equation as above. Detached cells were defined as single cells with no neighboring cells within 10 µm in distance. Highly invasive cells were defined as cells that invaded more than 200 µm from the channel’s edge. Cells with aspect ratio ≥ 2 were labeled ‘elongated’ and those with an aspect ratio < 2 were labeled ‘round’.

For invasion analysis of intracranial xenografts in athymic mice, fractal analysis was performed on 20X images of the tumor edge using the open-source imaging processing software Fiji (https://fiji.sc/). Images were first converted to grayscale using a built-in ImageJ function. Fractal analysis was performed using the Frac-Lac plug-in which utilizes a box-counting algorithm allowing for quantification of the morphological complexity of the tumor border. Four representative images were taken from each quadrant of the tumor edge to obtain an average value of the fractal dimension for each coronal section, with 3 sections imaged per tumor.

### Site directed biopsies

With UCSF IRB approval (#11-06160), biopsies were performed after obtaining informed consent utilizing a three-dimensional intraoperative navigation system for sampling the enhancing edge of the tumor and the central tumor core inside the enhancement before tumor resection commenced. Tumor samples were flash frozen in liquid nitrogen and stored at −80°C for further experimentation.

### CRISPR Knockout Screen and Next Generation Sequencing

The metabolism focused sgRNA libraries were designed and screen performed as previously described.^43,44^ Oligonucleotides for sgRNAs were synthesized by Genewiz and amplified by PCR. A total of 24,000,000 GBM43 cell line expressing sgRNAs were seeded into six 3D hydrogel invasion devices. After culture, invasive and non-invasive “core” cells were isolated by carefully disassembling the devices and isolating fractions by microdissection. Invasive cells were defined as those invading a distance greater than 200 µm from the channel wall. Genomic deoxyribonucleic acid (gDNA) was extracted using Monarch Genomic DNA Purification Kit (New England BioLabs, T3010S) following procedures for Tissue gDNA isolation. gDNAs from six invasion devices were pooled as one sample and amplified by PCR. PCR amplicons were then sequenced together with the initial and in vitro samples as per standard in vitro CRISPR-based screens. We then performed PCA analysis on normalized counts from each in vitro sample. Sequencing counts from samples were then summed, normalized (count per million), and analyzed as a single condition. The fitness score for each guide was calculated as log2 ratio of normalized counts. The median of the guides was used as the fitness score for each gene, and t test was used to assess whether the guides were significantly deviating from 0.

### Generating CRISPRi knockdown cells

A lentiviral plasmid containing a dCAS9/KRAB/MeCP2 cassette was obtained from Addgene. Lenti-X 293T cells were transfected using this plasmid and virus containing the plasmid was generated and appropriate titers were determined. GBM 43 cells were then transduced with the virus and selected using 5ug/ml blasticidin to obtain a pure dCAS9/KRAB/MeCP2 positive population. GBM43 cells expressing dCAS9/KRAB/MeCP2 were then transfected with the plasmid containing sgRNAs (**Supplementary Table 14**) and a puromycin resistance gene, followed by selection using 5 ug/ml Blasticidin and 5 ug/ml Puromycin. Gene knockdown was confirmed in all cases using qPCR before a functional invasion assay screen, with Western blot used for further validation of CTH knockdown before commencing additional studies of those cells.

### Metabolomics

Brain tumor tissue (4mg) was homogenized in extraction solution by mixing acetonitrile, isopropanol and water in proportions 3:3:2 (JT Baker, Center Valley PA), then vortexed for 45 seconds and then 5 minutes at 4C. Following centrifugation for 2 minutes at 14,000 rcf, two aliquots of the supernatant (500 μL each aliquot) were made for analysis and one for backup. One aliquot was dried via evaporation overnight in the Labconco Centrivap cold trap concentrator (Labconco, Kansas City MO). The dried aliquot was then resuspended with 500µL 50% acetonitrile (degassed as given), then centrifuged for 2 minutes at 14,000 rcf using the centrifuge Eppendorf 5415. The supernatant was moved to a new Eppendorf tube and again evaporated to dryness. Internal standards (C08-C30, fatty acid methyl esters) were then added and the sample was derivatized by methoxyamine hydrochloride in pyridine and subsequently by N-methyl-N-trimethylsilyltrifluoroacetamide for trimethylsilylation of acidic protons. Data were acquired as previously described.^45^ Briefly, metabolites were measured using a Restek corporation rtx5Sil-MS column (Restek Corporation; Bellefonte PA; 30 m length x 0.25 mm internal diameter with 0.25μm film made of 95% dimethyl/5%diphenylpolysiloxane) protected by a 10m long empty guard column which is cut by 20cm intervals whenever the reference mixture QC samples indicate problems caused by column contaminations. This sequence of column cuts has been validated by UC Davis Metabolomics Core with no detrimental effects detected with respect to peak shapes, absolute or relative metabolite retention times or reproducibility of quantifications. This chromatography method yields excellent retention and separation of primary metabolite classes (amino acids, hydroxyl acids, carbohydrates, sugar acids, sterols, aromatics, nucleosides, amines and miscellaneous compounds) with arrow peak widths of 2–3 seconds and very good within-series retention time reproducibility of better than 0.2s absolute deviation of retention times. The mobile phase consisted of helium, with a flow rate of 1 mL/min, and injection volume of 0.5 μL. The following mass spectrometry parameters were used: a Leco Pegasus IV mass spectrometer with unit mass resolution at 17 spectra s-1 from 80-500 Da at −70 eV for elution of metabolites. As a quality control, for each sequence of sample extractions, one blank negative control was performed by applying the total procedure (i.e. all materials and plastic ware) without biological sample. Result files were transformed by calculating the sum intensities of all structurally identified compounds for each sample (i.e. those signals that had been positively identified in the data pre-processing schema outlined above), and subsequently dividing all data associated with a sample by the corresponding metabolite sum. The resulting data were multiplied by a constant factor in order to obtain values without decimal places. Intensities of identified metabolites with more than one peak (e.g. for the syn- and anti-forms of methoximated reducing sugars) were summed to only one value in the transformed data set. The original nontransformed data set was retained. The general concept of this data transformation is to normalize data to the ‘total metabolite content’, but disregarding unknowns that might potentially comprise artifact peaks or chemical contaminants.

### Metabolic pathways analysis

Pathway enrichment analyses were performed for metabolites enriched in the invasive tumor front via MetaboAnalyst (version 5.0, www.metaboanalyst.ca)) to depict the most relevant metabolic pathways involving the identified features of the untargeted metabolomics dataset. The summary plot of the metabolite set enrichment analysis was implemented using hypergeometric testing to evaluate whether a particular metabolite set was represented more than expected by chance within the provided compound list. One-tailored p-values were provided after adjusting for multiple testing (Holm-Bonferroni method). The pathway analysis algorithm offers two parameters to determine relevant pathways within the comparison groups: statistical p-values derived from the quantitative enrichment analysis, and the pathway impact value calculated by the topological analysis with the relative-betweenness centrality.

### Lipidomics

Extraction of tumor tissue was carried out using a biphasic solvent system of cold methanol, methyl tertbutyl ether (MTBE), and water with some modifications. In more detail, 300 μL of cold methanol containing a mixture of odd chain and deuterated lipid internal standards [LPE(17:1), LPC(17:0), PC(12:0/13:0), PE(17:0/17:0), PG(17:0/17:0), d7-cholesterol, SM(d18:1/17:0), Cer(d18:1/17:0), sphingosine (d17:1), DG(12:0/12:0/0:0), DG(18:1/2:0/0:0), and d5-TG- (17:0/17:1/17:0)] was added to tissue aliquots in a 2 mL Eppendorf tube and then vortexed (10 s). Then, 1000 μL of cold MTBE containing CE 22:1 (internal standard) was added, followed by vortexing (10 s) and shaking (6 min) at 4 °C. Phase separation was induced by adding 250 μL of LC−MSgrade water followed by centrifugation at 14000 rpm for 2 min. The concentration of internal standards has been previously reported. Ten aliquots (each 100 μL) of the upper organic phase were collected and evaporated. The volumes of tissue samples and extraction solvents were used to ensure an aliquot for each platform and one backup. For a single platform, the method can be scaled as reported previously.^46,47^ Dried lipid extracts were resuspended using a methanol/toluene (9:1, v/v) mixture containing an internal standard CUDA (150 ng/mL), vortexed for (10 s), and centrifuged at 14000 rpm for 2 min prior to LC− MS analysis.

### RNA-sequencing

To extract RNA from invasion devices, devices were carefully disassembled, and the invasive cells were physically separated from the non-invasive “core” cells using microdissection. Samples were treated with 10k U/ml Hyaluronidase from bovine testes, Type IV-S (Sigma-Aldrich, H3884) for 15 min with agitation followed by 800 U/ml Proteinase K solution (Thermo Fisher Sci, AM2548) until the hydrogel was fully degraded. Afterwards, RNA was extracted using RNeasy Mini Kit (Qiagen, 74104) following manufacturer’s instructions. Invasive and Core RNA libraries were prepared and Illumina HiSeq NGS was performed (UC Davis Core; Davis, CA) per standard protocols. RNA-Seq raw files were processed via FASTQC for quality control followed by adaptor trimming using “cutadapt”. The files were then aligned by STAR alignment. The read counts were generated using the FeatureCounts. Differential gene expression analysis was carried out using the DESeq2 package in R.

#### Flash-frozen tumor samples

Brain tumor samples were homogenized in liquid nitrogen with mortar and pestle. Following homogenization, RNA was isolated from tissue using RNAeasy Mini Kit (Qiagen) following manufacturer’s instructions. Following extraction, RNA quantity an quality were assessed using NanoDrop™ 2000/2000c Spectrophotometers.

#### Online data set analysis

RNA-seq data for specific tumor anatomic structures in GBM, identified by H&E staining, was based on the Ivy Glioblastoma Atlas Project (Ivy GAP) (https://glioblastoma.alleninstitute.org/ and http://gliovis.bioinfo.cnio.es accessed on 27 Apr 2022).

### Nanostring multiplex transcriptomic analysis

RNA was isolated from paired site directed biopsies from the core and edge of three 3D invasion devices and three patient GBMs. A bioanalyzer was used to determine the quantity and quality of the RNA sample. RNA (100 ng) was used for the Metabolic Pathways Panel. RNA from each sample was hybridized with the codeset for 18 hours, and 30 μL of the reaction was loaded into the nCounter cartridge and run on the nCounter SPRINT Profiler. Enrichr software (Enrichr) was used to analyze the expression of pathways defined in the KEGG 2019 Human database and their statistical significance using the differentially expressed genes were obtained from the Nanostring nSolver software analysis as input. Pathway level expression changes for genes in the oxidative phosphorylation pathway (Kyoto Encyclopedia of Genes and Genomes entry HSA 00190) were visualized in R studio version 4.2.0 using the *pathview* function from the pathview package version 1.36.1. Briefly, the log2 fold change was computed for tumor invasive cells relative to tumor core cells averaged across bulk RNA-sequencing of three samples of invasive and core cells each. All differentially expressed genes were sorted in descending order by log2 fold change, and gene names were converted to ENTREZ IDs using utilities from the clusterProfiler package version 4.4.4. The upper and lower limits of the color scale were manually set to 2 and −2, respectively.

### qPCR

cDNA was created using qScript XLT cDNA Supermix (Quanta Bio), following the manufacturer’s protocol. cDNA was diluted to a constant concentration for all samples to ensure similar nucleic acid loading. Quantitative PCR was carried out using Power Syber Green Master Mix (Applied Biosystems), primer sequences in **Supplementary Table 15**, and an Applied Biosystems StepOne Real-Time PCR cycler following Applied Biosystems Syber guidelines: 95°C for 10 minutes, followed by 40 cycles of 95°C/15 sec and 60°C/1 min. Ct values were calculated using StepOne software (Applied Biosystems).

### Western Blots

Cell preparations were harvested in 1× radio immunoprecipitation buffer (10× RIPA; 9806, Cell Signaling) and one tablet each of PhoStop and Complete Mini (04906845001 and 04693124001, Roche). Insoluble materials were removed by centrifugation at 300 × g for 20 min at 4°C. Protein concentration was determined using the bicinchronic acid (BCA) assay (23225, Thermo Scientific). Samples were prepared with 30 μg of protein in RIPA buffer with 4× LDS loading buffer (LP0001, Life Technologies). Samples were electrophoresed on SDS/PAGE gels, transferred to PVDF membranes, and probed with anti-CTH (Cell Signaling #30068, 1:1000) and anti-GAPDH (Cell Signaling #5174, 1:50000) overnight at 4°C. Membranes were detected using anti-rabbit HRP-conjugated secondary antibodies (Cell Signaling #5127, 1:8000).

### ROS Measurements

ROS measurements in live cells were performed using Molecular Probes CellROX^TM^ Deep Red (Thermo Fisher Sci, C10422) or MitoSOX^TM^ Green (Thermo Fisher Sci, M36006) following manufacturer’s recommendations. For cells embedded in hydrogels, a concentration of 10 µM was utilized and probe incubation time was 30 minutes to enhance probe diffusion through hydrogel. Fluorescence imaging was performed on a Zeiss LSM 710 Laser Scanning Confocal Microscope. Immunostaining was performed to measure in-direct markers of ROS in frozen sections of hydrogel devices and patient tumors. Samples were probed with mouse anti-Nitrotyrosine (R&D Systems, MAB3248, 1:20) or mouse anti-Malondialdehyde (Thermo Fisher Sci, MA527559, 1:100) and later probed with goat anti-mouse Alexa Fluor 488 (Thermo Fisher Sci, A-11001, 1:200). Cell nuclei were visualized using 4′,6-diamidino-2-phenylindole dihydrochloride (DAPI, Sigma-Aldrich, 1 µg/ml).

### Cryosectioning

Prior to immunostaining, invasion devices were carefully disassembled. Hydrogels were fixed for 20 minutes in 4% PFA and rinsed twice with 1X PBS. Samples were cryoprotected by incubating hydrogels in 10% sucrose for 1 hour at room temperature followed by an overnight incubation in 30% sucrose at 4°C. Samples were equilibrated for 1 hour at room temperature in Tissue-Tek O.C.T. compound (Sakura Finetek USA, 4583) and then frozen at −80°C. Cryostar NX70 (Thermo Scientific) was used to make 10 µm-thick sections, which were immediately adhered to a Fisherbrand superfrost plus microscope slide (Thermo Fisher Sci, 22-037-246) and stored at −80°C until use. Samples were thawed for 10 min at room temperature, rinsed twice with 1X PBS, permeabilized with 0.05% Triton X-100 (Millipore Sigma, 9002-93-1) and blocked for 1 hour at room temperature with 5% Goat Serum in PBS prior to primary and secondary antibody incubation.

### Pharmacologic Inhibitor Assays

Small molecule inhibitors were reconstituted and handled according to manufacturer recommendations. For 2D cell viability assays, cells were seeded at 20k cells/ml in 48 well plates and incubated in a humidified 37°C chamber for 4-6 hours before replacing maintenance media with media containing pharmacologic inhibitors. After a 48-hour incubation, media and non-adherent cells were removed and rinsed with 1X PBS. Adherent cells were then fixed with 4% paraformaldehyde (PFA, Sigma-Aldrich) in 1X PBS for 15 min at room temperature and rinsed twice with 1X PBS. Cell nuclei were visualized using 4′,6-diamidino-2-phenylindole dihydrochloride (DAPI, Sigma-Aldrich, 1 µg/ml). Fluorescence imaging was performed on a Nikon Eclipse Ti-E epifluorescence microscope. Nuclei were counted using ImageJ and counts were normalized to that of control wells. Viability was confirmed for working concentrations using LIVE/DEAD Cell Imaging Kit (ThermoFisher) withthe highest dose that didn’t affect cell viability (**Supplementary Table 16**) after a 48-hour incubation used for invasion assays. All inhibitors and additional cell culture reagents (**Supplementary Table 17**) were replenished during media changes every 2 days, besides hydrogen peroxide which was incubated with cells for 30 minutes every 2 days.

### Murine Studies

Animal experiments were approved by the UCSF IACUC (approval #AN105170-02). 75,000 luciferase- and mCherry-expressing GBM43/dCAS9 or GBM43/CTHkd cells were implanted intracranially into the right frontal lobes of athymic mice (6-8 weeks, female) stereotactically. Mice for invasion fractal analysis were euthanized when BLI signal suggested tumor burden was approaching endpoint (2 to 4 weeks depending on the line), while a separate batch of tumors were taken to endpoint for Kaplan-Meier analysis.

## ACKNOWLEDGEMENTS

M.K.A. was supported by the NIH (R01CA227136, R01NS079697, and R01CA260443). J.H.G. was supported by an NCI diversity supplement awarded to 1R01CA227136. E.A.A was supported by the National Science Foundation (NSF) Graduate Research Fellowship. S.K. was supported by the NIH (R01CA227136, R01CA260443, and R01GM122375). K.J.W. was supported by NIH grant F31CA2228317. We thank Mary West of the Cell and Tissue Analysis Facility (CTAF) at UC Berkeley. This work was performed in part in the QB3 CTAF that provided the Cryostar NX70. Confocal imaging experiments were conducted at the CRL Molecular Imaging Center, RRID:SCR_017852, supported by the Gordon and Betty Moore Foundation. We would like to thank Holly Aaron and Feather Ives for their microscopy advice and support.

---

**Extended Data Figure 1.**
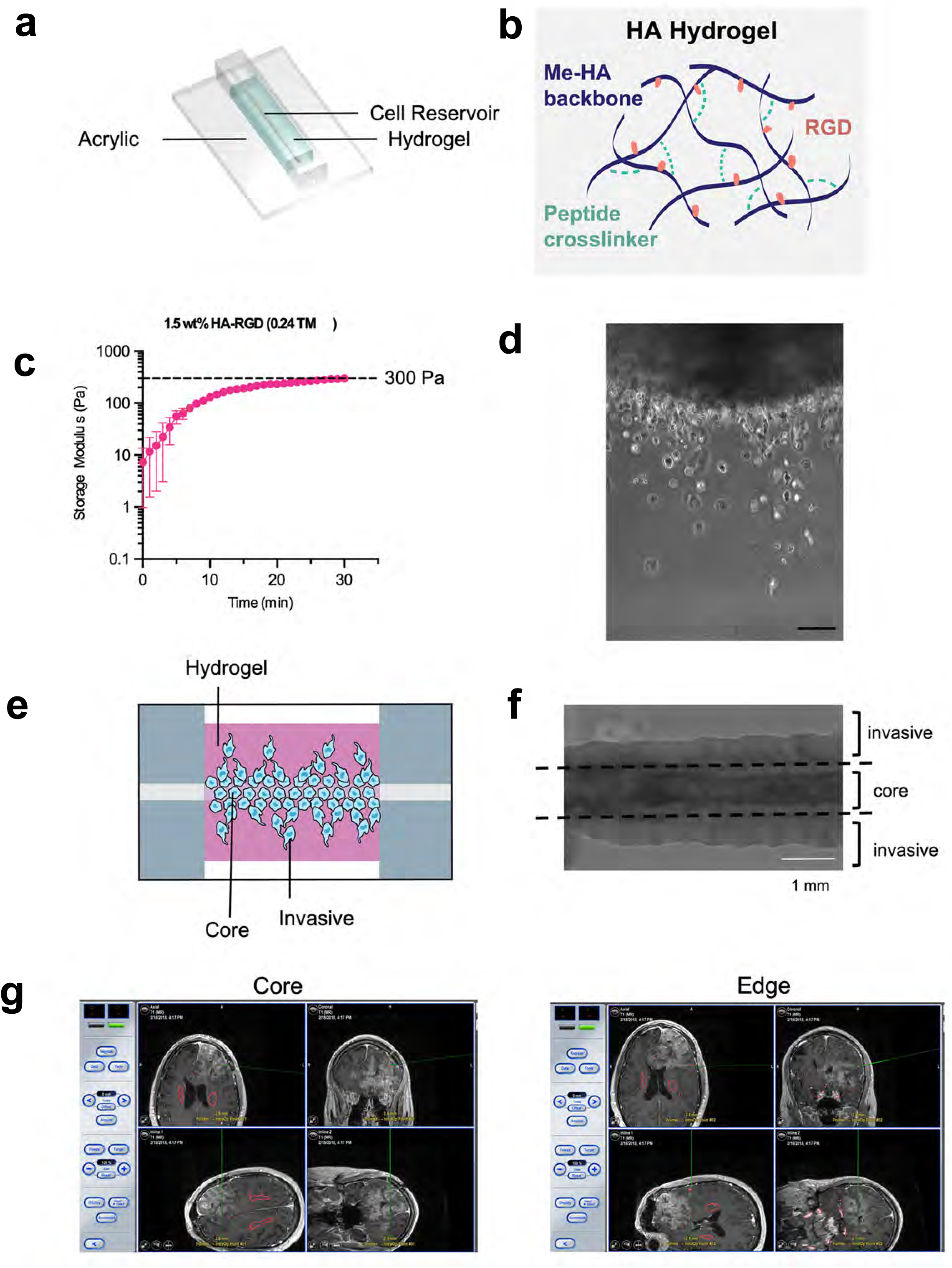
Setup for 3D hydrogel devices and patient site-directed biopsies. (**a**) Shown is an illustration of the invasion device which consists of a cell reservoir within a hyaluronic acid hydrogel on an acrylic slide. (**b**) HA-methacrylate (Me-HA) was functionalized with an integrin binding (RGD) peptide using the Michael-type addition reaction between the methacrylate groups on the polymer and the cysteine thiol groups on the peptide. The same addition reaction with the methacrylate groups was used to induce crosslinking with protease-sensitive peptide crosslinkers to form HA hydrogels. (**c**) Hydrogel storage modulus (G’) (in Pascals) versus time after the gel is casted. Time=0 represents the intersection of storage modulus (G’) and loss modulus (G’’) (n=3 hydrogels). (**d**) Phase image (10x magnification) of single cells of invasive fraction in HA hydrogel invasion device; scale bar=100 μm. (**e**) Illustration of invasion device (**f**) Phase image of invasion device with dotted lines indicating microdissection boundaries to isolate core and invasive cells. (**g**) Shown are representative screen shots taken from the neuro-navigation (BrainLAB system) illustrating locations of site-directed biopsies taken from the central core vs. enhancing edge of a newly diagnosed glioblastoma. The red voxels posterior to the gadolinium enhancement illustrate the bilateral locations of corticospinal tracts.

**Extended Data Figure 2.**
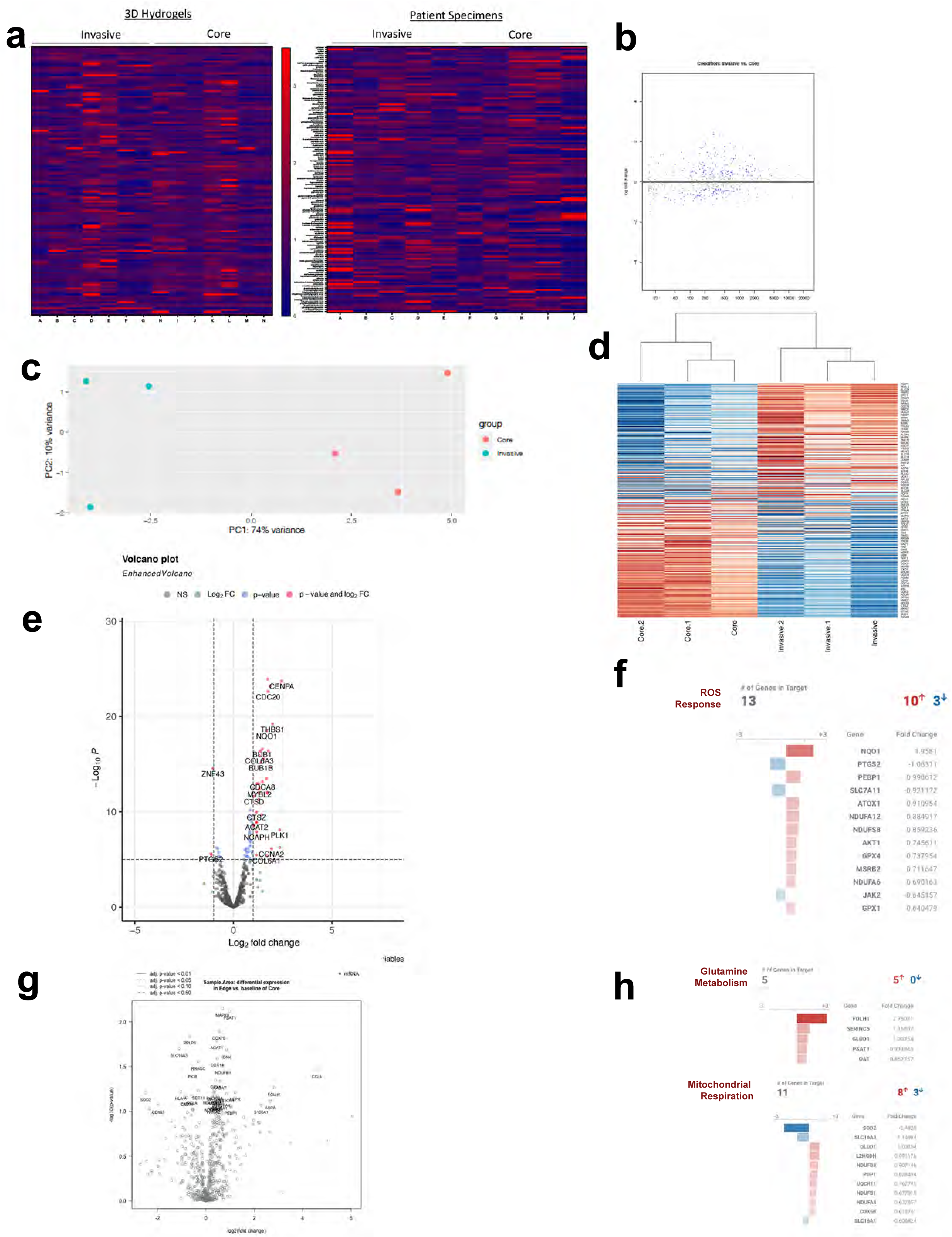
Metabolomic and transcriptomic analysis of invasive versus core GBM cells in 3D hydrogels and patient specimens. (**a**) Shown are levels of individual metabolites in the invasive fraction vs. core of 3D hydrogels after long-term culture (n=7 paired samples; left) and site-directed biopsies from patient tumor cores vs. invasive front (n=5 paired samples; right). (**b-e**) After GBM43 cells invaded 3D hydrogels during long-term culture, tumor cells in the invasive vs. core fractions were profiled (n=3 devices) using a 770 gene multiplex to analyze expression of metabolism genes, revealing: (**b**) an MA plot, in which the y-axis represents log_2_(fold change), the x-axis represents normalized RNA read counts of a particular gene, and each dot represents a gene, with < 0 indicating an increase in log_2_(fold change) in the Invasive fraction as compared to the Core fraction; (**c**) principal component analysis (PCA) of the data, showing that cells in the invasive front clustered together, but separate from cells in the tumor core; (**d**) a heatmap of the top 85 differentially expressed genes; (**e**) a Volcano plot illustrating differentially expressed genes in the invasive (right) and core (left) fractions; and (**f**) upregulated genes in the invasive fraction in the ROS response pathway. (**g-h**) Paired specimens from the invasive front vs. core of patient GBMs (n=3 pairs) were profiled using a 770 gene multiplex to analyze expression of metabolism genes, revealing (**g**) a Volcano plot illustrating differentially expressed genes in the invasive (right) and core (left) fractions; and (**h**) upregulated genes in the invasive fraction in the glutamine metabolism and mitochondrial respiration pathways.

**Extended Data Figure 3.**
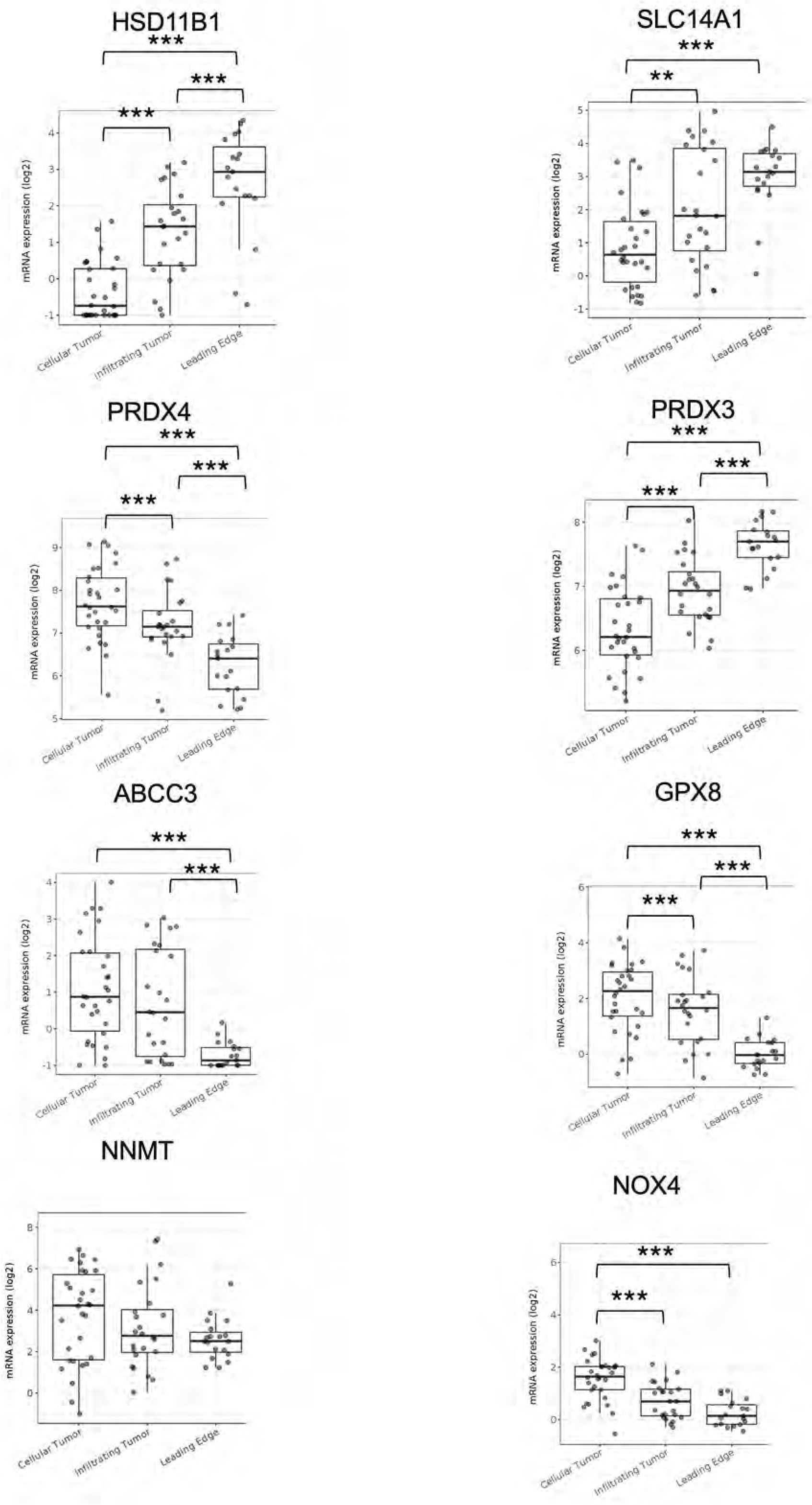
Validation of metabolic genes upregulated in the invasive front of patient GBM. Shown is the expression of 8 metabolic genes involved in the adaptive response to ROS that were found to be upregulated via RNA-seq in the invasive front versus core of patient GBMs. Validation was done using analysis of the Ivy Glioblastoma Atlas Project which sampled GBMs from 10 patients. *P< 0.05; **P< 0.01; ***P < 0.001.

**Extended Data Figure 4.**
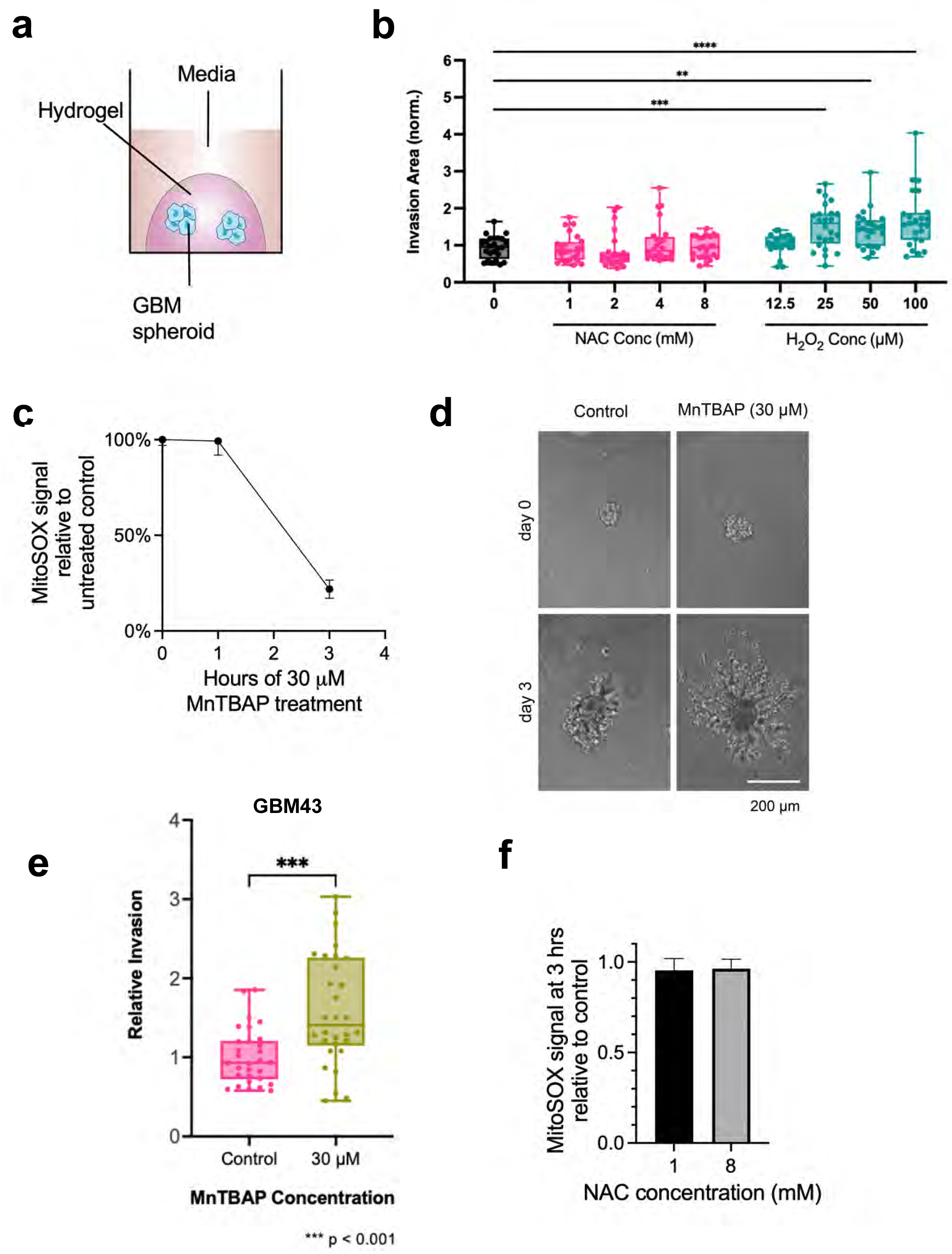
Hydrogen peroxide increases GBM invasion in 3D hydrogels. (**a**) Tumorsphere invasion assay schematic. **(b)** GBM43 spheroid invasion assay with various NAC and H_2_O_2_ concentrations (n=24 spheres, collected from 3 independent experiments). **(c)** Treating cultured GBM43 cells with 30 μM SOD mimetic MnTBAP lowered superoxide levels as assessed with the MitoSOX^TM^ dye by 80% within 3 hours. (**d**) 30 μM SOD mimetic MnTBAP increased invasion of GBM43 cells in tumorsphere invasion assay. **(e)** Quantification of data from (**d**). (**f**) NAC did not alter superoxide levels in GBM43 cells assessed by MitoSOX^TM^ (n=30 spheres, collected from 3 independent experiments). *P< 0.05; **P< 0.01; ***P<0.001; ****P<0.0001; (NAC=N-acetylcysteine).

**Extended Data Figure 5.**
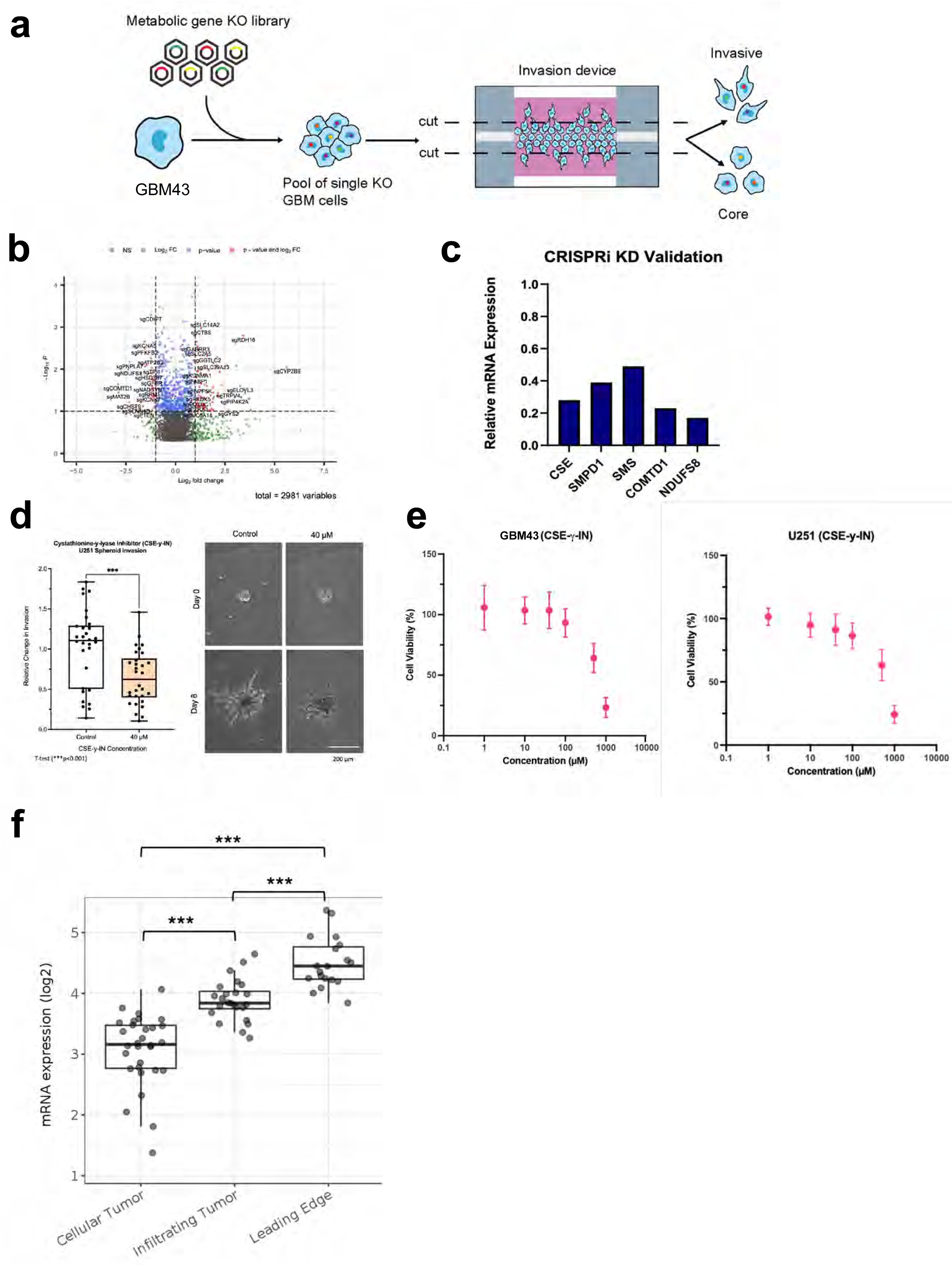
CRISPR knockdown screen of metabolic genes reveals a role for CTH in GBM invasion. (**a**) CRISPR Screening schematic. **(b)** Volcano plot displaying the enrichment of sgRNAs for metabolic genes in the core (left) and invasive front (right) of GBM 3D tumor models. (**c**) Results of qPCRs validating CRISPRi knockdown of five metabolic genes emerging as playing a role for GBM43 invasion into 3D hydrogels based on a CRISPRi screen of 3000 metabolic genes. (**d**) Results from tumorsphere invasion assays in U251 GBM cells treated with vehicle or 40 µM CSE-y-IN, a CTH small molecule inhibitor. (n=30 spheres, collected from 3 independent experiments *** P < 0.001). (**e**) Dose response curve for GBM43 (left) and U251 (right) GBM cells grown in varying concentrations of CSE-γ-IN for 48 hours after which cell viability was assessed (n=3 biological replicates). Based on these results, 40 μM of CSE-γ-IN was used for subsequent experiments. (**f**) Expression of pyridoxal kinase (PDXK), the enzyme that converts pyridoxine and other vitamin B6 precursors into pyridoxal-5′-phosphate (PLP), the bioactive form of CTH cofactor vitamin B6, in the invasive front versus core of patient GBMs was assessed using the Ivy Glioblastoma Atlas Project which sampled GBMs from 10 patients. *P< 0.05; **P< 0.01; ***P < 0.001; ****P<0.0001; CSE-γ-IN=Cystathioinine-γ-lyase inihibitor.

**Extended Data Figure 6.**
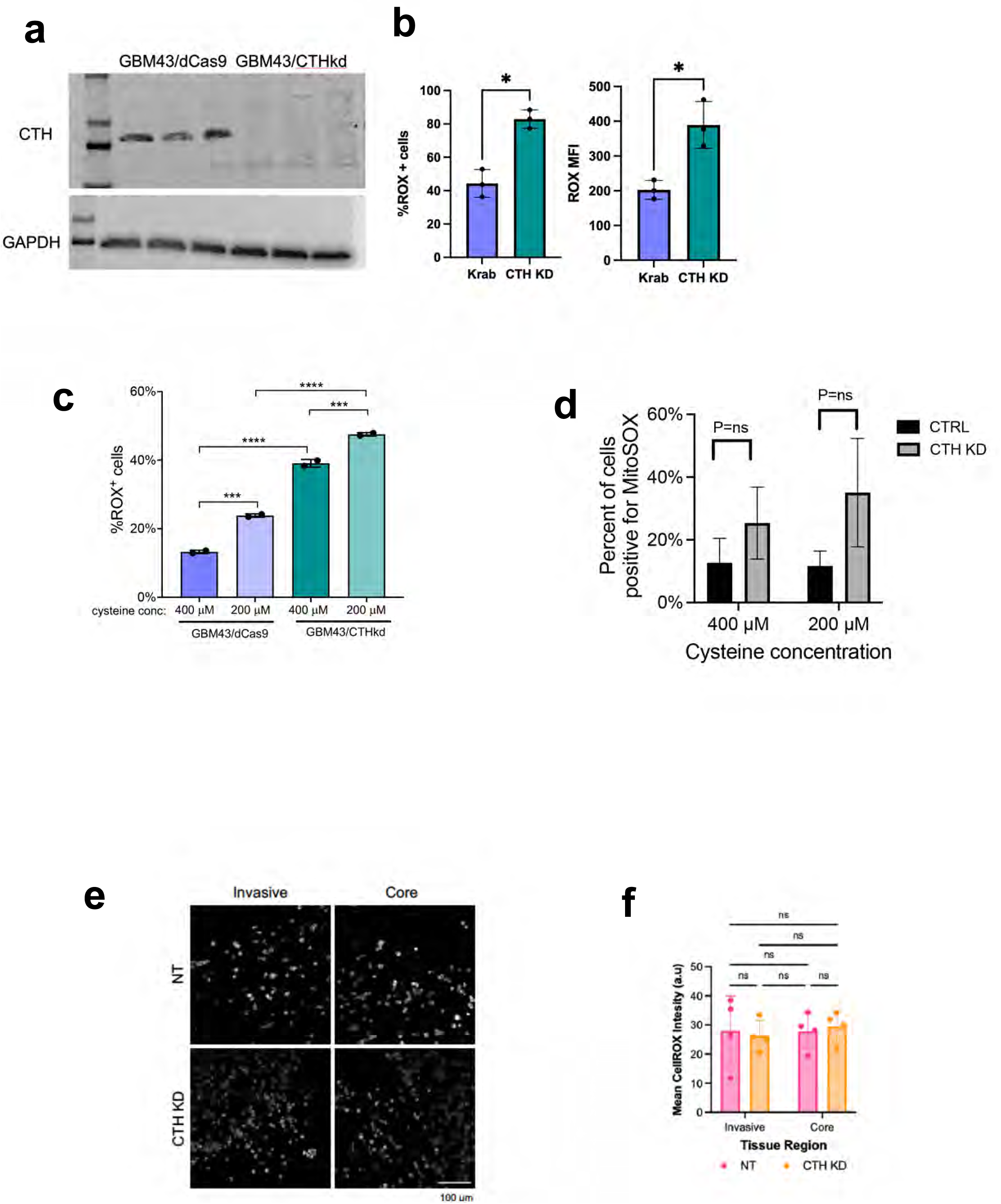
Effect of CTH knockdown on GBM cellular ROS levels during invasion. (**a**) Western blot validating CRISPRi knockdown of CTH in GBM43 cells (three replicates of GBM43/dCas9 in the left most lanes, three replicates of GBM43/CTHkd in the right most lanes). (**b**) CTH knockdown increases ROS levels in GBM43 cells as determined using the CellROX reagent and measuring percent of cells that are CellROX positive (left; P=0.01) or mean fluorescence intensity (MFI; P=0.03; right). (**c**) Cysteine deprivation increased ROS accumulation in GBM43 cells expressing dCas9-KRAB with or without sgRNAs targeting CTH. (**d**) CTH knockdown did not affect superoxide levels in GBM43 cells in 400 or 200 μM cysteine, as determined using the MitoSOX reagent and measuring percent of cells that are MitoSOX positive. (**e-f**) Cellular ROS levels were measured by the CellROX reagent in GBM43 control and CTH KD cells isolated from 3D invasion devices, by (**e**) imaging the cells incubated with CellROX after completing the invasion assay and (**f**) measuring mean fluorescence intensity (P=ns, n=4 devices). *P< 0.05; **P< 0.01; ***P < 0.001; ****P<0.0001

**Extended Data Figure 7.**
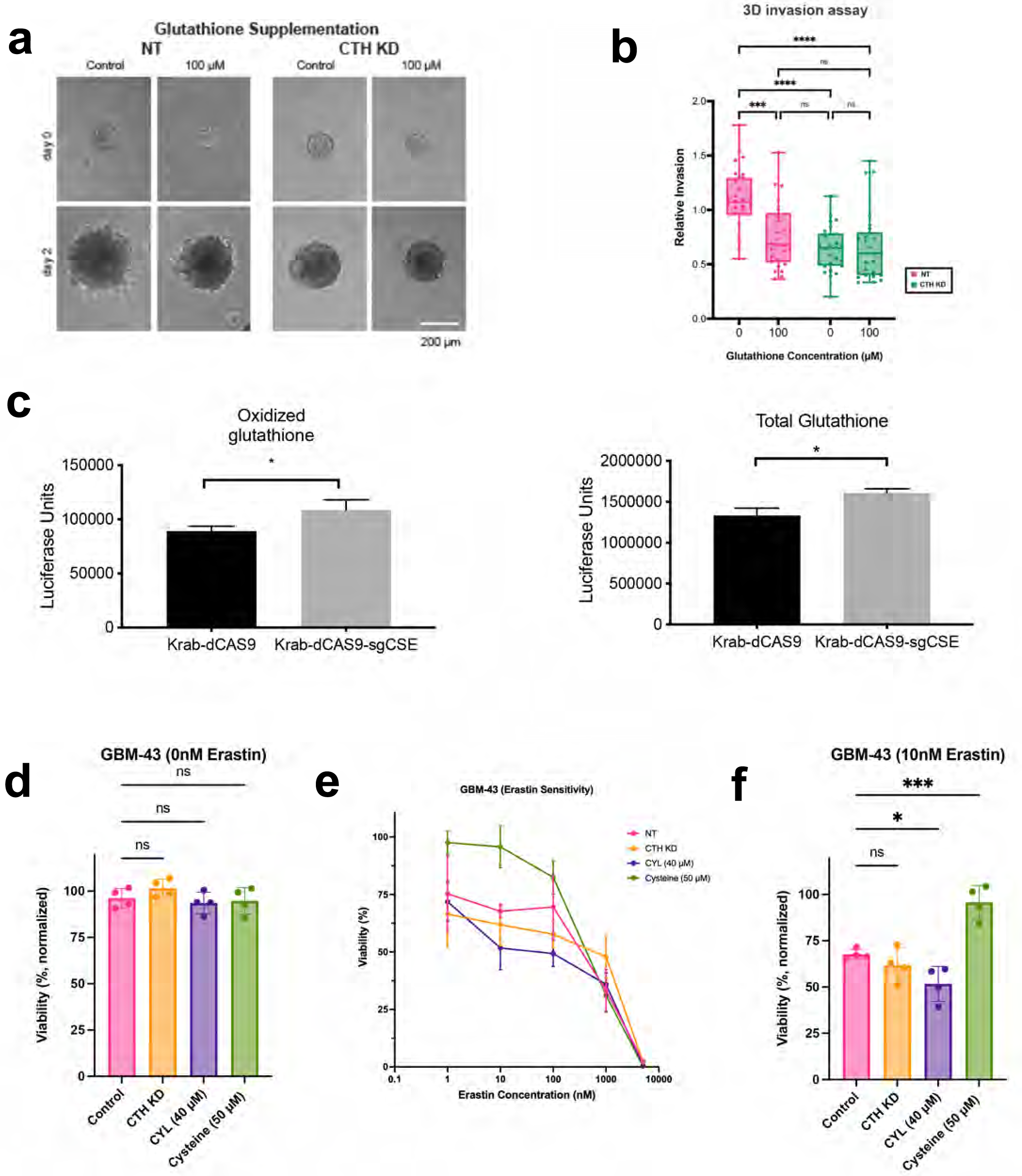
CTH drives GBM invasion due to cysteine not glutathione biosynthesis. (**a-b**) The impact of glutathione supplementation on invasiveness of GBM43 cells with CTH knockdown was assessed through **(a)** images of GBM43 control and CTH KD spheroid invasion assay in 3D Hydrogels with glutathione supplementation, from which (**b**) invasion was quantified and normalized relative to control cells without glutathione (n=26 spheres, collected from 3 independent experiments). (**c**) When GBM43 cells expressing Krab-dCAS9 and sgRNA targeting CTH were compared to GBM43 cells expressing just Krab-dCAS9, oxidized (left; P=0.03) and total (right; P=0.01) glutathione levels were slightly elevated. (**d-e**) The effect of CTH knockdown, treatment with 40 µM CSE-y-IN, or 50 µM additional cysteine (total cysteine=450 µM) was assessed on the response of GBM43 cells to ferroptosis inducer erastin (n=4 biological replicates). (**d**) Baseline cell viability was not affected by CTH knockdown, treatment with 40 µM CSE-y-IN, or 50 µM cysteine (P=ns). (**e**) Cell viability in 10 nM erastin was reduced by 40 µM CSE-y-IN (P<0.05) and increased by 50 µM cysteine (P<0.001). *P< 0.05; **P< 0.01; ***P < 0.001; ****P<0.0001.

**Extended Data Figure 8.**
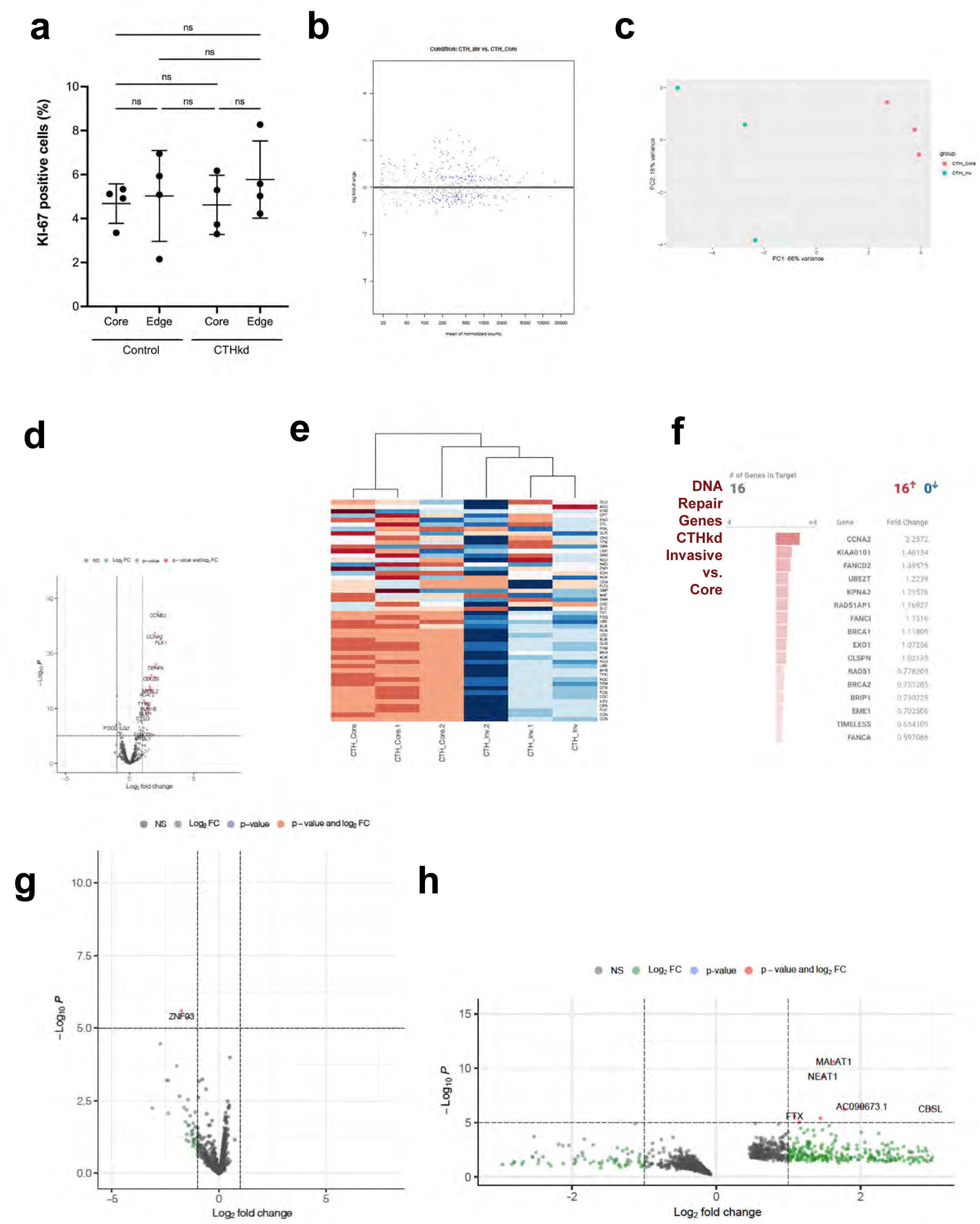
CTH drives GBM invasion due to cysteine not glutathione biosynthesis. (**a**) KI-67 staining of dCas9 and CTHkd GBM43 cells in hydrogel invasion devices revealed no differences in proliferation between core vs. edge fractions or between control and knockdown cells (n=4 devices). (**b-f**) GBM43 cells with CRISPRi targeting of CTH were seeded into invasion devices, after which cells isolated from core and invasive fractions were transcriptomically assessed using the NanoString nCounter platform and the 770 metabolic gene multiplex, revealing (**b**) an MA plot, in which the y-axis represents log_2_(fold change) and the x-axis represents normalized RNA read counts of a particular gene, with each dot representing a gene, and < 0 indicating an increase in log_2_(fold change) in the Invasive fraction as compared to the Core fraction; (**c**) PCA plots in which cells in the invasive front clustered together and apart from cells in the core; (d) Volcano plot from CTHkd core (log_2_fold change<0) versus CTHkd invasive (log_2_fold change>0) comparison; (**e**) heatmap identified enriched metabolic genes in the invasive fractions relative to the core fractions of the hydrogels; and (**f**) GSEA identified several upregulated pathways in CTHkd cells in the invasive fraction versus the core fraction, including the DNA repair genes listed here. (**g**) Upregulated genes from the 770 metabolic gene multiplex in GBM43 cells with CTH knockdown in the invasive fraction were similar to genes upregulated in control GBM43 cells in the invasive fraction, creating a Volcano plot with log2fold change>0 representing genes upregulated in invasive CTH kd cells relative to invasive control GBM43 cells and log2 fold change<0 representing genes upregulated in invasive control GBM43 cells relative to invasive CTH kd cells, revealing few differentially expressed genes between the invasive fractions of the two cell types. (**h**) Bulk RNA-seq was used to compare invasive versus core CTH KD cells, creating a Volcano plot, with log2fold change>0 representing genes upregulated in invasive cells and log2 fold change<0 representing genes upregulated in core cells.

**Extended Data Figure 9.**
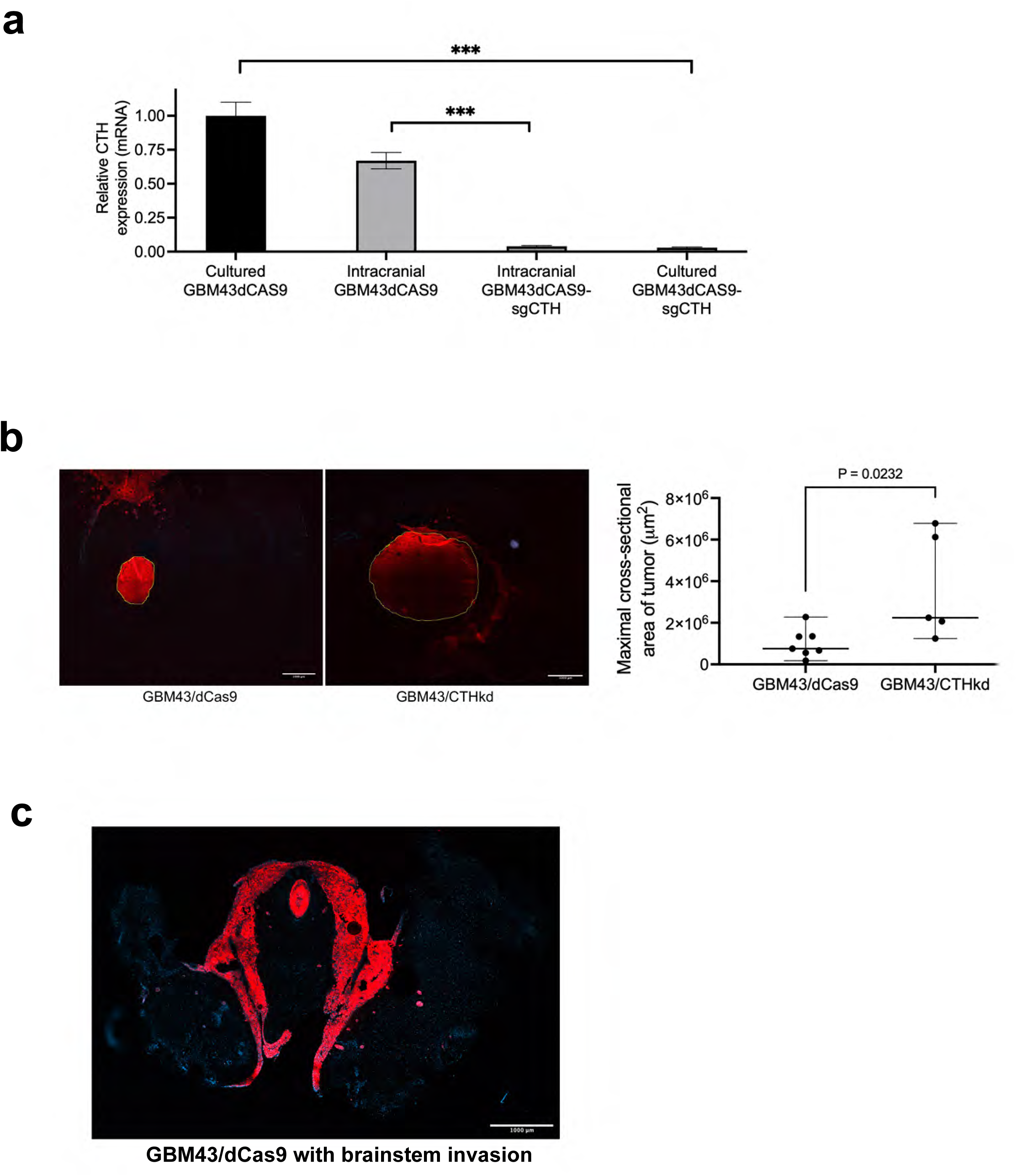
CTH knockdown slows GBM43 invasion *in vivo*. (**a**) After mice carrying intracranial GBM43 xenografts with CRISPRi targeting of CTH reached endpoint, the tumors maintained their knockdown of CTH, as assessed by qPCR of explanted tumors. (**b-c**) Compared to intracranial control GBM43 xenografts, GBM43 xenografts with CRISPRi targeting of CTH (**b**) had a larger maximal cross-sectional area when assessed just before endpoint (P=0.02), but were (**c**) less invasive, as measured in **Fig. 6f** and exemplified here by brainstem invasion seen in 25% of intracranial GBM43/dCas9 xenografts, but not seen in GBM43/CTHkd xenografts.

