## Supplementary Table 16 for "Multi-omic screening of invasive GBM cells in engineered biomaterials and patient biopsies reveals targetable transsulfuration pathway alterations"

| **Pharmacologic Inhibitor** | **Target** | **Concentration** |
| --- | --- | --- |
| Metformin (Cayman Chemical) | Complex 1 | 500 µM |
| Deguelin (Cayman Chemical) | Complex 1 | 0.5 µM |
| Cystathionine-y-lyase-IN-1 (Med Chem Express) | CTH (CSE) | 10 – 40 µM |
| Entacapone (Sigma) | COMT | 0.5 µM |
| Desipramine (Sigma) | SMPD1 (ASM) | 0.5 µM |
| 5'-Deoxy-5'-methylthioadenosine (Cayman Chemical) | SMS (MTA) | 50 µM |

**Supplementary Table 16. Drug concentrations used for invasion assays.** Shown are the concentrations used and vendors for the six drugs used to target the products of the metabolic genes that emerged from a CRISPR screen of metabolic genes whose targeting slowed GBM invasion in hydrogel devices. These concentrations represent the highest concentrations which did not affect GBM43 cell viability after 48 hours in culture.
