## Supplementary Table 17 for "Multi-omic screening of invasive GBM cells in engineered biomaterials and patient biopsies reveals targetable transsulfuration pathway alterations"

| **Additional Cell Culture Reagents** | **Concentration** |
| --- | --- |
| N-Acetyl-L-cysteine (Sigma-Aldrich, A9165) | 1 – 8 mM |
| L-Glutathione (Sigma-Aldrich, G6013) | 100 µM |
| MnTBAP (Sigma-Aldrich, 475870) | 30 µM |
| L-Cysteine (Sigma-Aldrich, C7477-25G) | 50 µM |
| Erastin (MedChemExpress, HY-15763) | 1 nM – 5 µM |
| Hydrogen peroxide solution (Sigma-Aldrich, H1009) | 12.5 – 100 µM |

**Supplementary Table 17. Concentrations used for additional cell culture reagents.** Shown are the concentrations used and vendors for reagents used for cell culture experiments.
